# Comparative high-throughput phenotyping across two facilities reveals differential impact of defence mechanisms on plant growth and development

**DOI:** 10.64898/2026.03.20.713143

**Authors:** Sylvain Poque, Murilo Sandroni, Pedro Caparros, Jesper Cairo Westergaard, Katriina Mouhu, Manzur E Mohsina Ferdous, Erik Andreasson, Laura J Grenville-Briggs, Åsa Lankinen, Thomas Roitsch, Kristiina Himanen, Erik Alexandersson

## Abstract

Fitness costs of plant disease defence are often subtle and difficult to quantify. In this study, we therefore used comparative high-throughput phenotyping in two independent facilities to assess growth, morphology and physiology of potato (cv. Désirée) with high time-resolution monitoring different defence mechanisms under pathogen-free conditions. Plants were either treated weekly with the resistance inducers β-aminobutyric acid (BABA; 10 mM) or potassium phosphite (KPhi; 36 mM) or comprised six transgenic lines expressing late blight resistance genes (single Rpi genes or a three-gene stack) or reduced jasmonate perception (StCOI1-RNAi).

Over four weeks, image-derived traits revealed consistent cross-facility effects for plant height and colour: BABA treatment increased plant height but reduced canopy area and induced a paler greenness signature, whereas KPhi caused minimal and transient growth effects. Chlorophyll fluorescence at the NaPPI facility indicated reduced vitality (Rfd_Lss) in BABA-treated plants and increased Rfd_Lss following KPhi, while maximum PSII efficiency was largely unchanged. Several transgenic lines showed somewhat reduced above-ground biomass. Enzyme activity profiling produced distinct treatment and genotype signatures, but was strongly modulated by facility conditions that overrode these specificities. Overall, high-throughput phenotyping robustly detected subtle growth–defence trade-offs across platforms.

**Highlight:** High-throughput optical phenotyping validated across two independent research facilities reveals that stacked resistance genes and resistance inducers in potato trigger subtle growth trade-offs.

**Graphical abstracts:** 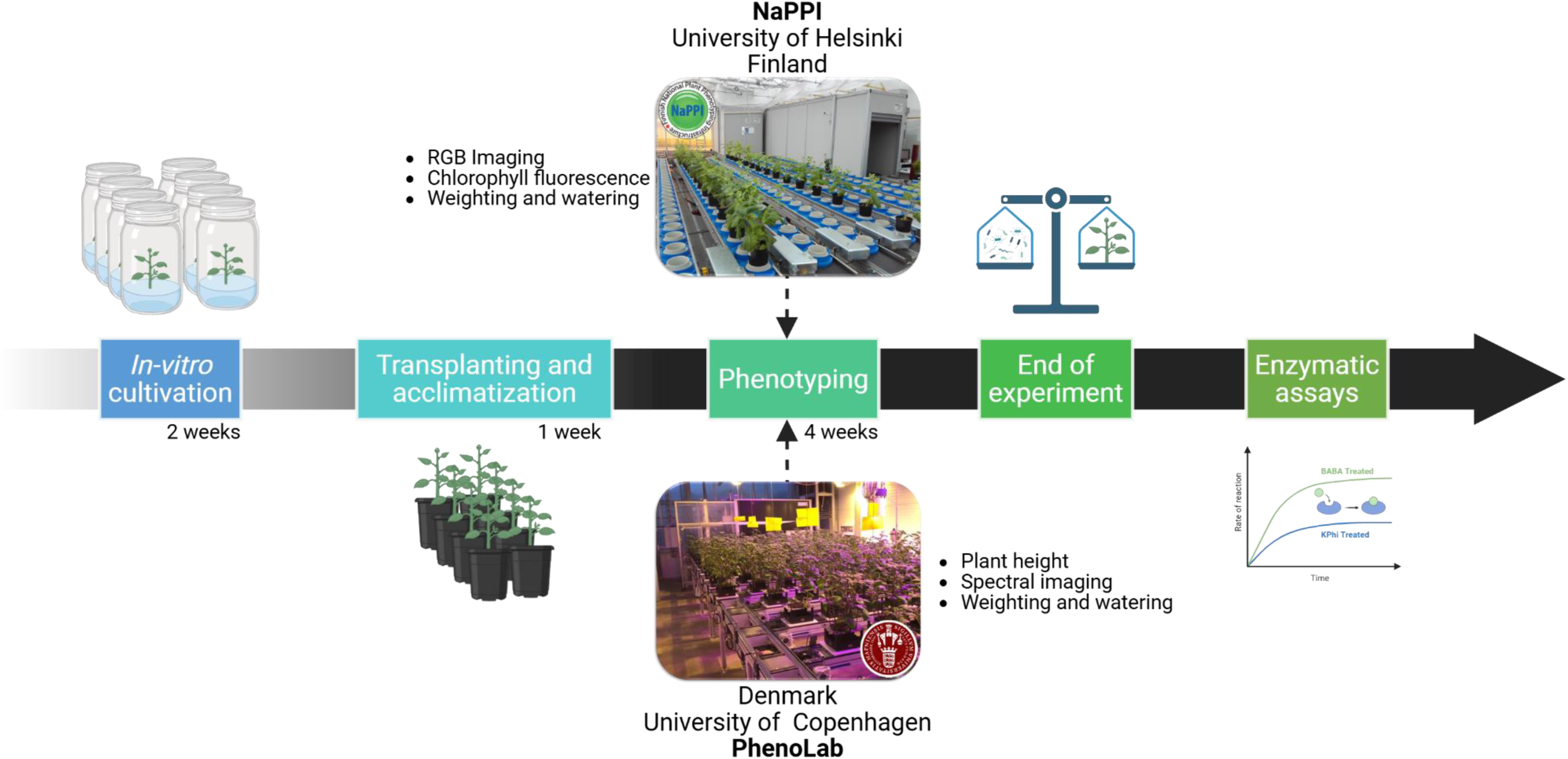

Experimental timeline for high-throughput plant phenotyping platforms.

Created in BioRender. Poque, S. (2026) https://BioRender.com/nmkve7g

## Introduction

High-throughput plant phenotyping in controlled climates represents an ideal tool to objectively monitor plant performance and quantify defence-related costs. Yet, the intersection between the studies of biostimulants and plant phenotyping is surprisingly underexplored **(De Diego and Spíchal, 2022),** and costs related to plant defence mechanisms even less so. Plants can employ different defence strategies against pests and pathogens, ranging from metabolically costly constitutive defences that are always active to more energy-efficient induced defences triggered only upon threat detection **(Vos *et al*., 2013; Karasov *et al*., 2017)**. For example, cell surface-localized pattern-recognition receptors (PRRs) recognise conserved pathogen-associated molecular patterns (PAMPs), and intracellular nucleotide-binding domain leucine-rich repeat containing receptors (NLRs; also termed Resistance or R genes) recognise pathogen effectors **(Wang *et al*., 2023)**. PRRs activate PAMP-triggered immunity (PTI) and NLRs initiate effector-triggered immunity (ETI) **(Bigeard *et al*., 2015)**. PTI and ETI are interdependent, activating common signalling components and capable of potentiating each other to strengthen the overall plant immune response **(Yuan *et al*., 2021; Ngou *et al*., 2022)** and can both trigger systemic responses in the plant, known as systemic acquired resistance (SAR) **(Vlot *et al*., 2021)**. Plants can also respond to soil microbes, leading to induced systemic resistance (ISR), or to exogenous compounds sometimes referred as resistance inducers (RIs) **(De Kesel *et al*., 2021)**, of which β-aminobutyric acid (BABA) is extensively studied **(Cohen *et al*., 2016)**. SAR is generally SA-dependent, while ISR is microbe-induced and primarily regulated by JA and ethylene, often after root perception.

Studies of costs of resistance to pathogens over the past 50 years have revealed that trade-offs between defence and growth are often caused by pleiotrophic gene effects and antagonistic cross-talk among plant hormones **(Brown and Rant, 2013; Karasov *et al*., 2017; He *et al*., 2022)**. A study in tomato and potato found that only a small subset of R genes was highly expressed during infection **(von Dahlen *et al*., 2023)**, indicating that fitness costs may mainly result from downstream signalling rather than R gene expression itself.

Detailed molecular understanding regarding costs of defence is of great importance for agriculture, e.g. in relation to plant breeding for resistant cultivars **(Zhang *et al*., 2023)**, including pyramiding of R genes or alternatively the use of Susceptibility (S-genes) to avoid resistance breakage **(Garcia-Ruiz *et al*., 2021)**. However, fitness costs associated with disease resistance can be difficult to quantify without large sample sizes or very accurate measures of lines in constant environmental conditions **(Cipollini *et al*., 2017)**. Significant advances have been made in furthering our understanding of the mechanisms behind IR, but a better understanding of the associated costs of defence in plants is necessary if IR will have a broader implementation in agriculture **(Sandroni *et al*., 2020)**. Induced resistance can be triggered by the application of chemical compounds including non-protein amino acids, such as β-amino butyric acid (BABA) and inorganic salts, i.e. potassium phosphite (KPhi). Studies in Arabidopsis have shown that chemical induction of resistance by BABA, even in the absence of pathogen infection, can reduce growth and seed production, particularly at higher doses **(Van Hulten *et al*., 2006)**. However, it has been hypothesised that such fitness costs may be greater in model plants than in crops, as crop breeding may have selected alleles that confer defence benefits at lower cost. Therefore, it is important to assess resistance costs in crops to ensure that mechanisms like IR can be used for protection without reducing yield. Notably, KPhi and BABA, tested here, have also been reported to act as biostimulants, improving yield and quality in various crop species **(Calvo *et al*., 2014; Trejo-Téllez and Gómez-Merino, 2018)**.

In potato, both BABA and KPhi can control late blight caused by the oomycete pathogen *P. infestans* to a certain extent, but also other diseases **(Liljeroth *et al*., 2010, 2016; Alexandersson *et al*., 2016)**. BABA has been shown to reduce late blight lesions by up to 50% in greenhouse grown plants **(Liljeroth *et al*., 2010)**. KPhi provides field protection against late blight in starch potato cultivars, matching full fungicide applications in resistant varieties and offering full protection when combined with half-dose fungicide **(Liljeroth *et al*., 2016; Mulugeta *et al*., 2019)**. Both BABA and KPhi activate defence-related genes in plants resulting in an IR phenotype **(Burra *et al*., 2014; Lankinen *et al*., 2016)**. Furthermore, KPhi application has been shown to alter the abundance of metabolites **(Wu *et al*., 2019)**, and BABA also affects both the transcriptome and the proteome **(Bengtsson *et al*., 2014b)**.

It is vital not only to test the efficiency of various plant defence strategies but also to build knowledge and capacity to systematically study possible fitness costs related to these defence strategies. Increased resistance is typically associated with growth and fitness penalties; however, the extent of these costs differs depend on the type of resistance mechanism involved **(Cipollini *et al*., 2017).**

Costs trade-offs can arise from the allocation of resources to defence and the activation of direct or indirect defence responses. Accordingly, source/sink relations and defence mechanisms were shown to be co-ordinately regulated in response to pathogen related stimuli **(Ehness *et al*., 1997)** or pathogen infection **(Berger *et al*., 2007)**. Thus, cell physiological phenotyping via profiling of activity signatures of key enzymes of carbohydrate and antioxidant metabolism **(Jammer *et al*., 2022)** has been proven as valuable approach to characterize the impact of pathogen infection **(Pandey *et al*., 2021; Spanic *et al*., 2023)** and are expected to be also proxy for the cost of defence **(Großkinsky *et al*., 2015)**.

To better understand the potential fitness costs of resistance in an agriculturally relevant crop plant, we conducted experiments in two Nordic plant phenotyping facilities: the PhenoLab at University of Copenhagen and the National Plant Phenotyping Infrastructure at the University of Helsinki **(Alexandersson *et al*., 2018)**. Comparing plant phenotypic traits across independent phenotyping facilities is essential for evaluating the reproducibility and methodological robustness of high-throughput phenotyping systems and also testing potential subtle fitness costs as those related to plant defence activation. Nevertheless, yet there are very few studies done under these experimental conditions. Differences in environmental control, sensor configuration, imaging protocols, and data processing pipelines can introduce facility-specific biases that affect trait estimation. Cross-facility trait measurements enable systematic benchmarking, support the harmonization of phenotyping protocols, and help distinguish technical variability from biologically meaningful signals. Such methodological validation is critical for ensuring the comparability and broader utility of phenotypic data generated across platforms. It can also pinpoint more subtle differences as for growth-induction of defence responses studied here.

By comparative high-throughput phenotyping across two facilities, our overall aim was to evaluate whether heightened plant defence in potato results in fitness costs without pathogen pressure. To study the effect of one R-gene vs. R-genes pyramiding (the stacking of multiple resistance genes in one genotype), we selected four transgenic lines of the potato cultivar Désirée, where three have broad spectrum resistance to *P. infestans* **(Ghislain *et al*., 2019)**. To test whether fitness costs would be incurred when jasmonic acid hormone sensing was disrupted, we used two lines in which the expression of the Coi1 gene (a jasmonate receptor) was knocked down using RNAi. To understand how IR could affect plant development, the IR-compounds BABA and KPhi were used to induce resistance in potato plants. With the use of image-based sensors, we measured growth in area and plant height, morphological traits and colour during four weeks of growth. To assess physiological costs, we measured PSII maximum quantum yield as a proxy measure for photosynthetic activity. We also collected end point growth measurements including shoot and tuber weight as well as activity of key enzymes of carbohydrate and antioxidant metabolism. These data allowed us to visualise small variations in plant growth, development and physiology and suggest that the pyramiding of Rpi genes or the chemical-induction of IR in potato may confer subtle allocation costs in the absence of disease.

## Material and methods

### Plant material

*Solanum tuberosum* cultivar Désirée (wild type, WT) was selected for its moderate susceptibility to *P. infestans*, along with six transgenic potato lines generated in the Desirée Holland background (Table 1): two lines carrying a combination of three Rpi genes, [Rpi-blb1, Rpi-blb2 and Rpi-vnt1.1, named 3R1 and 3R2 **(Ghislain *et al*., 2019; Wang *et al*., 2020)**], R2A (over-expressing the R2 resistance gene **(Lenman *et al*., 2016)**), blb1 [over-expressing the broad spectrum late blight resistance gene Rpi-blb1 **(Van Der Vossen *et al*., 2003)**], and two independent coronatine insensitive StCOI1-RNAi plants (named coi1H1 and coi1X5) with reduced sensitivity to jasmonic acid methyl ester **(Halim *et al*., 2009)**. To test resistance inducers (RI), Désirée (cv.) was used for the treatments with BABA and KPhi. In all experiments six biological replicates were used both for the lines and for the RI treatments. The same Désirée (cv.) plants were used as control for both sets of plants.

**Table 1.** Plant material grown at the National Plant Phenotyping Infrastructure (NaPPI) and PhenoLab.

| Genotype | Late blight resistance | Reference |
| --- | --- | --- |
| ***Cultivar*** |  |  |
| **Désirée** | Moderately susceptible | **(Andrivon *et al.*, 2007)** |
| ***Transgenic lines in Désirée (cv.)*** |  |  |
| **3R1** | Expression of RB and Rpi-blb2 from *Solanum bulbocastanum*<br>Rpi-vnt1.1 from *Solanum venturii* | **(Wang *et al.*, 2020)** |
| **3R2** | Expression of RB and Rpi-blb2 from *Solanum bulbocastanum*<br>Rpi-vnt1.1 from *Solanum venturii* | **(Wang *et al.*, 2020)** |
| **coi1H1** | Reduced sensitivity to jasmonic acid methyl ester | **(Halim *et al.*, 2009)** |
| **coi1X5** | Reduced sensitivity to jasmonic acid methyl ester | **(Halim *et al.*, 2009)** |
| **R2A** | Expression of a R2 resistance gene Rpi-ABPT | **(Lenman *et al.*, 2016)** |
| **blb1** | Expression of the broad-spectrum resistance gene Rpi-blb1 | **(Van Der Vossen *et al.*, 2003)** |

### Plant growth conditions

The experiments were conducted in two high-throughput phenotyping platforms (HTPPs) to take advantage of the different sensors in each location and for the comparability of some measured traits that were common in both sites. Both HTPPs consist of an automatic conveyor system for the movement of the pots in combination with a phenotyping station with specific sensors for plant monitoring and measurements.

The first experiment was conducted at the Modular Plantscreen™ Conveyor System (Photon Systems Instruments, Drásov, Czech Republic) of the National Plant Phenotyping Infrastructure (NaPPI) of the University of Helsinki Vikki campus during the spring of 2019. *In-vitro* plantlets were generated from cuttings at SLU, Alnarp, Sweden. Plants were grown under sterile conditions for two weeks in tissue culture in a Phyto chamber with a day length of 16 h (light conditions: 140 µmol m^-2^ s^-1^) at a temperature of 22 °C. The plantlets were then transported to NaPPI in the *in-vitro* containers and upon arrival were transplanted firstly to small pots of 8 cm diameter for *ex-vitro* acclimation. After 18 days, the seedlings were transferred to 5 L plastic pots filled with “Kekkilä Karkea Ruukutusseos W R8014” (Kekkilä, Finland) and brought to the Modular Plantscreen™ conveyor belt system of the facility for automated watering and imaging for four weeks. Automated watering was controlled using pot-weighing, allowing daily recording of pot weights and maintaining each plant at a target total weight of 3.3 kg. The plants were fertigated once with 2 µS Kekkilä Vegetable-Suberex solution (Kekkilä, Finland) at the beginning of the experiment.

The environment was programmed to an 18-h photoperiod with targeted temperatures of 20 °C during the day and 18 °C at night. Environmental parameters were recorded at one-minute intervals and included air temperature, relative humidity (RH), and light intensity (LI). Based on the recorded data, the actual mean daytime temperature was 25.6 °C, with a maximum of 35.9 °C and a minimum of 18.7 °C. During the night, the mean temperature was 21.1 °C, with maximum and minimum values of 30.1 °C and 18.6 °C, respectively. Mean daytime RH was 37.5%, while nighttime averaged 48.8%. When natural light intensity fell below the target threshold, supplemental illumination was supplied using high-pressure sodium (HPS) lamps, resulting in a mean daytime light intensity of 248.8 µmol m⁻² s⁻¹. Daytime light intensity ranged from 11.0 to 1868.0 µmol m⁻² s⁻¹.

For the second experiment in the fall of 2019, the *in-vitro* plantlets were transplanted into 1.5 L plastic pots filled with a commercial pot soil (Yrkesplantjord, Weibulls, Sweden) and grown in a controlled environment chamber at Biotron, SLU Alnarp, at 20 °C, 65 % RH, and a photoperiod of 16 h of light (160 µmol m^-2^ s^-1^) for one week before they were transferred to the fully automated plant phenotyping facility PhenoLab at the University of Copenhagen Taastrup campus **(Amby *et al*., 2025)**. The seedlings were transplanted to square pots, 13 x 13 cm, filled with 2,2 L of commercial substrate (Krukväxtjord Lera & Kisel, Hammenhög, Sweden) with the addition of PG-Mix powder fertilizer. The seedlings were grown with natural greenhouse light and supplemented lighting to provide a 16-h photoperiod by usage of white tubular Osram LED lamps giving light intensity (at 150 MM) 120 µmol m^-2^ s^-1^with day/night temperatures maintained at 20/18°C with 70% of relative humidity. In addition, automated watering and image sessions in the morning and afternoon, as well as randomization were carried out every 6 hours for five weeks. Environmental parameters were recorded at three-minute intervals and included air temperature, relative humidity (RH), and light intensity (LI). Based on the recorded data, the actual mean daytime temperature was 22.8 °C, with a maximum of 25.1 °C and a minimum of 17.9 °C. During the night, the mean temperature was 18.8 °C, with maximum and minimum values of 24.3 °C and 17.4 °C, respectively. Mean daytime RH was 56.93%, while nighttime averaged 57.3%. Illumination was provided using LED lamps, resulting in a mean daytime light intensity of 224.3 µmol m⁻² s⁻¹. Daytime light intensity ranged from 27.4 to 472.3 µmol m⁻² s⁻¹.

### Induced resistance (IR) treatments

The IR treatments started seven days after plantlets were transferred to the two-phenotyping facilities and were carried out on a weekly basis. Plants were sprayed once a week on four occasions, with 10 mL of tap water (control), 10 mL of BABA 10 mM (DL-3-aminobutyric acid, Sigma A44207) or 10 mL of 1.25% (v/v) Proalexin (LMI AB, Helsingborg, Sweden), which corresponds to 36 mM of potassium phosphite (KPhi).

### RGB image acquisition and trait extraction

For the first experiment conducted in NaPPI, the visible light Red Green Blue (RGB) top (RGB2) and side view (RGB1) imaging was performed every second day for top view and once a week for side view during the experiment to collect data on growth in area and height, morphological traits and colour. Side- and top-down view images were captured using an RGB line scan and top-down cameras (IDS Imaging Development Systems GmbH, Obersulm, Germany), respectively. Top view RGB images of the plants were captured at resolution 2560 × 1920 pixels and camera height was automatically adjusted by light curtain unit measuring plant height. For side view images, line scan camera captured images at resolution of 2560 × 3476 pixels. The RGB images of each individual plant were processed by first removing the background using pixel-based colour thresholding, followed by the extraction of morphological parameters and colour-based segmentation using the MorphoAnalysis v. 1.0.5.1 software (Photon Systems Instruments, Drásov, Czech Republic). During the experiment, plant height was determined using two methods: weekly extraction from the side-view images (RGB1) and daily measurements from a light curtain. For the RGB1 measurement, three side-view images were taken at each time point, with a 120° rotation of the plant between each imaging. Plant height was then calculated as the average of values obtained from the three angles. Plant canopy area and greenness were quantified from top-view images (RGB2) with GREENNESS parameter computed using Python version 3.10.7 and OpenCV **(Bradski, 2000)**. For each image, the pixel values of the Red (R), Green (G), and Blue (B) channels were averaged to obtain mean values (avgR, avgG, avgB). GREENNESS parameter was then calculated using the formula described by Signorelli et al. (2023):

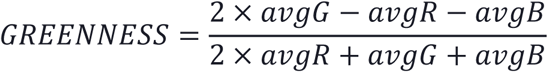

### Biomass measurements at the end of the experiment

At harvest (approximately 38 days after transplanting, DAT), plants were weighed for aboveground fresh matter (FM, in g), and tubers were counted (number of tubers, NT) and weighed (tuber weight, TubW, in g). Plant shoots were bagged and dried at 75°C for three days, after which above ground dry matter (DM, in g) was recorded.

### Photosystem II photochemistry measurements

Chlorophyll fluorescence (ChlF) measurements were carried out in the NaPPI facility using a top-view Pulse Amplitude Modulated (PAM) chlorophyll a fluorometer (FluorCam FC-800MF, PSI, Czech Republic) equipped with a dark-adaptation tunnel **(Tschiersch *et al*., 2017)**. ChlF imaging was used to monitor plant physiological responses and detect stress indicators at the end of the experiment. ChlF parameters were measured after a 15-min dark-adaptation period using a quenching protocol developed in FluorCam 7.0 software (PSI, Drásov, Czech Republic). The quenching protocol comprised two phases: a dark-adapted phase followed by a light-adapted (actinic) phase. In the dark-adapted phase, minimal fluorescence (F_0_) was recorded and a saturating pulse was applied to obtain maximal fluorescence (F_m_); variable fluorescence was determined as F_v_=F_m_-F_0_, and the maximum PSII quantum yield as F_v_/F_m_. In the subsequent light-adapted phase, actinic light was applied, allowing measurement of F_p_, the fluorescence level immediately after light onset, while saturating pulses were used to determine maximal fluorescence in the light (F_m__Lss) and the steady-state fluorescence (F_t__Lss). The effective PSII quantum yield at light steady state was calculated as QY_Lss=(F_m__Lss-F_t__Lss)/F_m__Lss. To evaluate non-photochemical dissipation under actinic light, steady state nonphotochemical quenching was computed as NPQ_Lss=(F_m_-F_m__Lss)/F_m__Lss and the ratio of fluorescence decline at light steady state as Rfd_Lss=(F_p_-F_t__Lss)/F_t__Lss. A higher Rfd_Lss indicates a stronger decline from initial to steady-state fluorescence, reflecting greater activation of dissipative processes; importantly, the related Rfd is widely used as a vitality index as it correlates with photosynthetic performance and overall plant condition **(Haitz and Lichtenthaler, 1988; Lichtenthaler *et al*., 2005)**.

### Multispectral imaging

For the second experiment conducted in PhenoLab, imaging was performed using a multispectral system capturing ten discrete spectral bands from UVA to NIR. All images were acquired from the top view only. The system used a JAI BM 500GE camera equipped with a Sony ICX625 2/3″ CCD monochrome sensor (2456 × 2058 pixels). Plants were illuminated sequentially in ten narrow spectral bands, each 10–20 nm wide, covering the range of 365–970 nm. This setup enabled the acquisition of calibrated reflectance data across ten image layers for each plant as it passed through the imaging station.

The acquisition mode alternated between crop coverage (CC) priority and reflectance priority, with the priority mode typically switched at midday on weekdays. Imaging was scheduled twice daily for four weeks, and the plants were randomized in the greenhouse after each imaging session. Throughout the trial, quality control images were collected to detect overexposure and other imaging artefacts that required correction.

Post-trial, all multispectral images were processed in VideometerLab version 3.12.25 (Videometer, Herlev, Denmark) to generate segmentation algorithms isolating plant tissue **(Pandey *et al*., 2025).** These segmentations were used to extract crop coverage and band-specific reflectance values. NDVI was calculated from pixel-wise reflectance values, with avgNIR computed as the average reflectance of the three NIR channels (780, 890, and 970 nm) and avgRED as the average reflectance of the red channels (645 and 670 nm):

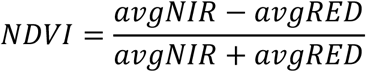

To enable comparison with NaPPI-derived data, plant greenness in PhenoLab was computed using the same GREENNESS metric described above, where avgG corresponds to the green 525 nm band, avgB to the blue 460 nm band, and avgR to the average of the two red bands at 645 and 670 nm.

Due to the physical constraints of the PhenoLab system, approximately 60 cm in height clearance and 40 cm in width, the crop coverage (CC) imaging, which is always performed with the camera positioned 60 cm above the pot surface, is only relevant for part of the growth period. Plant height was measured twice daily using a line scan light curtain with 2D snapshot technology (Model VSPM-6F2413, AG, Waldkirch, Germany) positioned before entry into the vision chamber.

### Enzymatic assays

At the end of both experiments, four to six leaflets were sampled from the top of the plant (third to sixth leaf) and quickly frozen in liquid nitrogen. Samples were kept in -80 ℃ until further processing. For the enzymatic assays, frozen potato leaves were homogenized in liquid nitrogen and extraction was done according to the protocol reported by **Jammer et al. (2015)**. A total of 500 mg of ground plant material from each sample was mixed with 1 mL of extraction buffer (40 mM TRIS-HCl pH 7.6, 3 mM MgCl_2_, 1 mM EDTA, 0.1 mM PMSF, 1 mM benzamidine, 14 mM β-mercaptoethanol, 24 μM NADP) for 60 minutes at 4 °C. The homogenate was centrifuged at 4 °C and 20,000 *g* for 45 min. The supernatant (crude extract) was aliquoted to new tubes and kept on ice. The pellet was washed three times with distilled water, re-suspended in 1 mL of high salt buffer (1 M NaCl, 40 mM TRIS-HCl pH 7.6, 3 mM MgCl_2_, 1 mM EDTA, 0.1 mM PMSF, 1 mM benzamidine, 14 mM β-mercaptoethanol, 24 μM NADP), and mixed by continuous shaking at 4 °C overnight. The homogenate was centrifuged at 4 °C and 20,000 *g* for 25 min and the supernatant (cell wall extract) was transferred to new tubes. Both the cell wall extract and the crude extract were dialysed overnight with 20 mM potassium phosphate buffer (pH 7.4) at 4 °C. The extracts were aliquoted, frozen in liquid nitrogen, and stored at −20°C until further use.

For the estimation of carbohydrate enzyme activities, the method of **Jammer et al. (2015)** was used, while antioxidant enzyme activities were measured following the method of **Fimognari et al. (2020)**. A 96-well microtiter plate (Sarstedt, Nümbrecht, Germany) was used for the semi-high-throughput analysis of kinetic assay of C enzymes: phosphoglucose isomerase (PGI, EC5.3.1.9), phosphoglucomutase (PGM, EC 2.7.5.1), UDP-glucose pyrophosphorylase (UGPase, EC 2.7.7.9), ADP-glucose pyrophosphorylase (AGPase, EC 2.7.7.27), glucose-6-phosphate dehydrogenase (G6PDH, EC 1.1.1.49), sucrose synthase (Susy, EC 2.4.1.13), aldolase (Ald, EC4.1.2.13), cell wall invertases (cwInv, EC 3.2.1.26), vacuolar invertase (vacInv, EC 3.2.1.26) and cytosolic invertase (cytInv, EC 3.2.1.26) and A enzymes: glutathione reductase (GR, EC 1.8.1.7), monodehydroascorbate reductase (MDHAR, EC 1.6.5.4), catalase (CAT, EC 1.11.1.6), dehydroascorbate reductase (DHAR, EC 1.8.5.1), glutathione peroxidase (GPX, EC 1.11.1.9), guaiacol peroxidase (POX, EC 1.11.1.7), ascorbate peroxidase (APX, EC 1.11.1.11), and apoplastic class III peroxidases (aPOX, EC 1.11.1.7) with a 2–3 μL protein extract. For each sample determinations were done with three technical replicates and mean values are shown for the biological replicates,

### Data processing for detecting outliers

Time series phenotypic data were cleaned using the statgenHTP **(Millet *et al*., 2025)** package in R (version 4.3.3, https://www.R-project.org/). Outlier detection followed the package’s built-in workflow, which evaluates the consistency of each observation relative to neighbouring within the plant level trajectory. The procedure applies local regression to approximate each time series using parametric functions. A local regression model is first fitted at selected anchor points along the time course and then interpolated across intermediate points to obtain a smooth expected trajectory. Confidence intervals around the fitted values are then computed, and any observations falling outside these intervals are flagged as outliers. Data points identified as outliers were subsequently removed prior to all downstream analyses.

### Principal Component Analysis

PCA were performed using Python (version 3.13, http://www.python.org) on standardized enzyme measurements. Prior to analysis, rows containing missing values were removed to ensure a complete numeric matrix. All enzyme variables were z-scored using the StandardScaler transformer from scikit-learn **(Pedregosa *et al*., 2012)**. A two-component PCA model was then fitted to the scaled data. Sample scores for the first two principal components (PC1 and PC2) were used to visualize treatment-dependent separation among samples, while loading vectors were overlaid to show the contribution and direction of each enzyme in the biplot. All data handling and preprocessing were performed using pandas **(McKinney, 2010)**, and PCA biplot visualizations, were generated using matplotlib **(Hunter, 2007)** and seaborn **(Waskom, 2021)**.

### Statistical analysis

Generalized additive mixed models (GAMMs) were used to analyse time-series data, capturing non-linear temporal trends and group-specific trajectories. Time (days after transplanting, DAT) was modelled with penalized splines, with smooth terms varying by IR treatment or transgenic line to allow distinct temporal responses across groups. To account for the experimental design, we included time-varying individual-plant effects using factor-smooth interactions. In the NaPPI facility, block (Block 1: 25 plants; Block 2: 23 plants) was included as a factor-smooth interaction term to account for spatial growth differences arising from plant placement within the greenhouse. Because residuals in repeated-measures data are typically autocorrelated, we incorporated an AR(1) correlation structure and fitted all models using REML with automatic smoothness selection in the R package mgcv **(Wood, 2000)**. Model adequacy was evaluated using standard GAM diagnostics and inspection of residual autocorrelation. Fitted smooths and pairwise smooth differences, comparing transgenic lines to the Desiree line, or IR treatments to untreated Desiree, were visualized using the R package itsadug **(van Rij et al., 2025)**.

Single timepoint traits, dry and fresh matter, tuber number and weight, as well as ChlF parameters for transgenic lines from the NaPPI facility were analyzed using a linear mixed effects model (R package lme4,**Bates et al. (2015)**) with Genotype as a fixed effect and Block as a random intercept to capture greenhouse placement–related growth differences. For other datasets (e.g., NaPPI IR and all PhenoLab), where a block structure was absent, a fixed effects linear model from base R’s stats package was used to estimate genotype or treatment effects for all single-timepoint traits. For all analyses, estimated marginal means (EMMs) followed by Dunnett-type contrasts were used to compare each transgenic line or IR treatment with its designated control, with Bonferroni adjustment for multiple comparisons (R package emmeans, **Lenth and Piaskowski (2017)**). All plotting were performed with R package ggplot2 **(Hadley Wickham, 2016)**.

## Results

### Induced resistance (IR) treatments

We evaluated potential costs associated with applying plant resistance inducers β-aminobutyric acid (BABA) and potassium phosphite (KPhi) to plants grown in two distinct high-throughput phenotyping (HTPP) facilities. Plants were assessed for growth, development, and physiological traits over a period of four weeks.

### Plant height following IR treatments over the course of the experiment

Daily plant height measurements throughout the experiment showed that BABA-treated plants grew taller than KPhi-treated plants and control plants in both the NaPPI and the PhenoLab facilities (Fig. 1). In contrast, KPhi-treated plants did not differ significantly from the control. Together, these results demonstrate treatment reproducibility across both facilities.

**Fig. 1.**
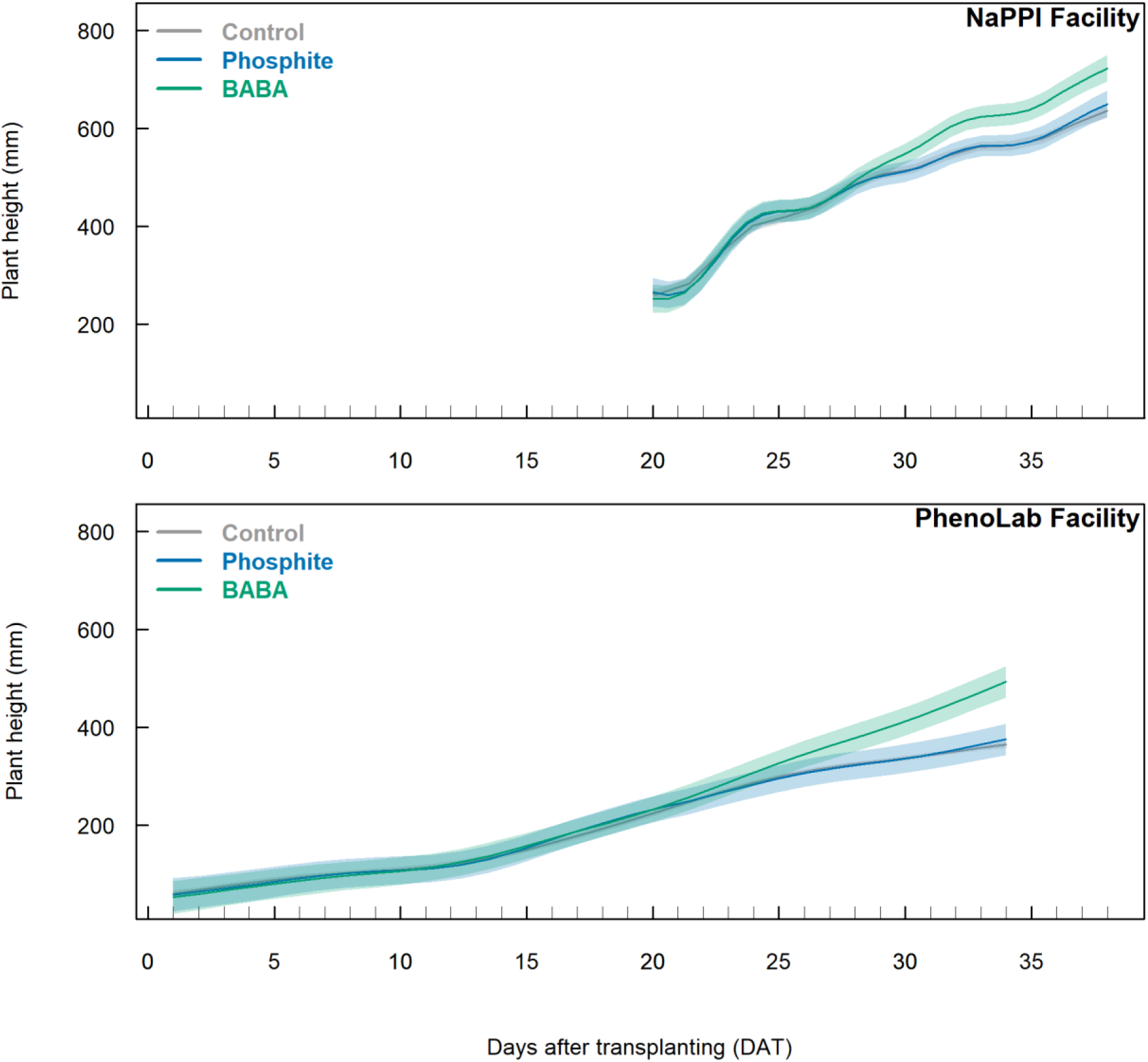
Plant height dynamics in response to induced-resistance treatments. Generalized Additive Mixed Models (GAMMs) were used to analyse plant height measured in NaPPI (upper panel) and PhenoLab (lower panel). Group differences were evaluated using two-sided pairwise smooth-difference tests derived from the GAMMs, with significance concluded when the 95% confidence interval of the difference did not include zero. Six biological replicates per treatment were used in each facility, and IR treatments were applied weekly via foliar sprays. Control plants are represented in grey, BABA-treated plants in blue, and phosphite-treated plants in green. Shaded bands represent the 95% confidence intervals of the GAMM fits and indicate periods of significant differences.

### Extracted plant canopy traits following IR treatments over the course of the experiment

Total plant canopy area extracted from the top view images over the whole growth in the NaPPI facility (Fig. 2A) provided additional information about the growth of the plants. To assess treatment effects relative to the control, pointwise estimated differences were calculated, and treated conditions were considered to significantly diverge from the control when their 95% confidence intervals did not include zero. At the beginning of the experiment, BABA and KPhi-treated plants showed no significant differences in canopy area compared to the control (Figs. 2B and 2C). However, from 20 to 35 DAT the canopy area of BABA-treated plants was significantly smaller than that of the control (Fig. 2C). In contrast, KPhi-treated plants had a significantly larger canopy area than the control during the mid-stage of growth (from 21 to 25 DAT; Fig. 2B), followed by a smaller canopy area toward the end of the time course, with significance observed briefly from 32 to 35 DAT and again from 37 to 38 DAT (Fig. 2B).

**Fig. 2.**
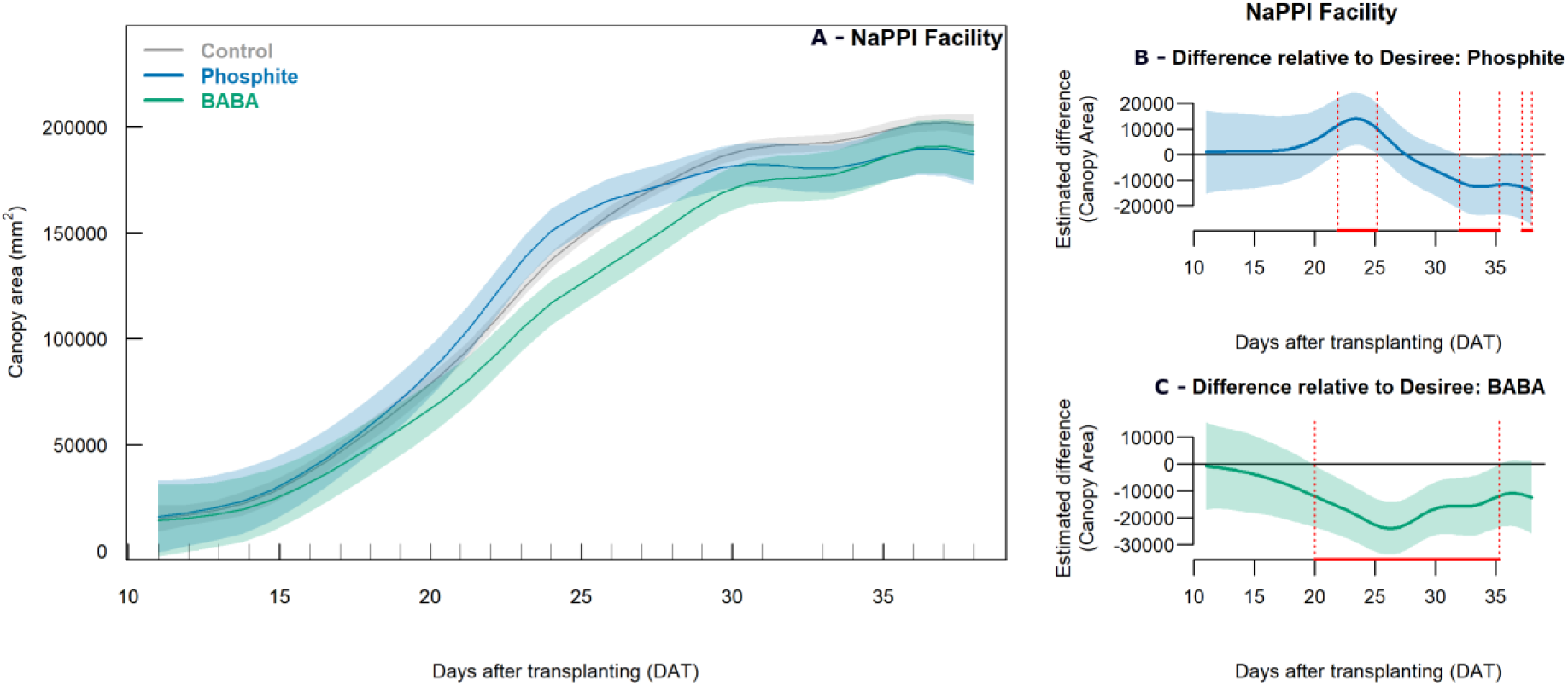
Canopy area dynamics in response to induced-resistance treatments. Generalized Additive Mixed Models (GAMMs) were used to analyse canopy area measured in NaPPI (A). Pointwise differences between the fitted canopy area of the Desiree (cv.) control and phosphite treated plants (B), or BABA treated plants (C), are shown across days after transplanting (DAT). Group differences were evaluated using two sided pairwise smooth difference tests derived from the GAMMs, with significance concluded when the 95% confidence interval of the difference did not include zero. Six biological replicates per treatment were used, and IR treatments were applied weekly via foliar sprays. Control plants are shown in grey, BABA treated plants in green, and phosphite treated plants in blue. Shaded bands represent the 95% confidence intervals of the GAM fits. Red lines at the bottom of panels B and C indicate periods during which significant differences at the 95% confidence level were detected.

### Colour composition following IR treatments over the course of the experiment

Top-view RGB images from NaPPI and single-channel Red, Green, and Blue images from PhenoLab were used to calculate the greenness index, enabling analysis of potential stress responses to IR treatments through color composition. Greenness profiles of BABA-treated plants were consistently and significantly higher than those of the control from 24 to 38 DAT in NaPPI (Fig. 3B) and from 20 to 34 DAT in PhenoLab (Fig. 3D), whereas KPhi-treated plants remained similar to the control in both facilities (Figs. 3A and 3C). This increase in greenness for BABA-treated plants reflects an overall paler green color compared to the control (data not shown).

**Fig. 3.**
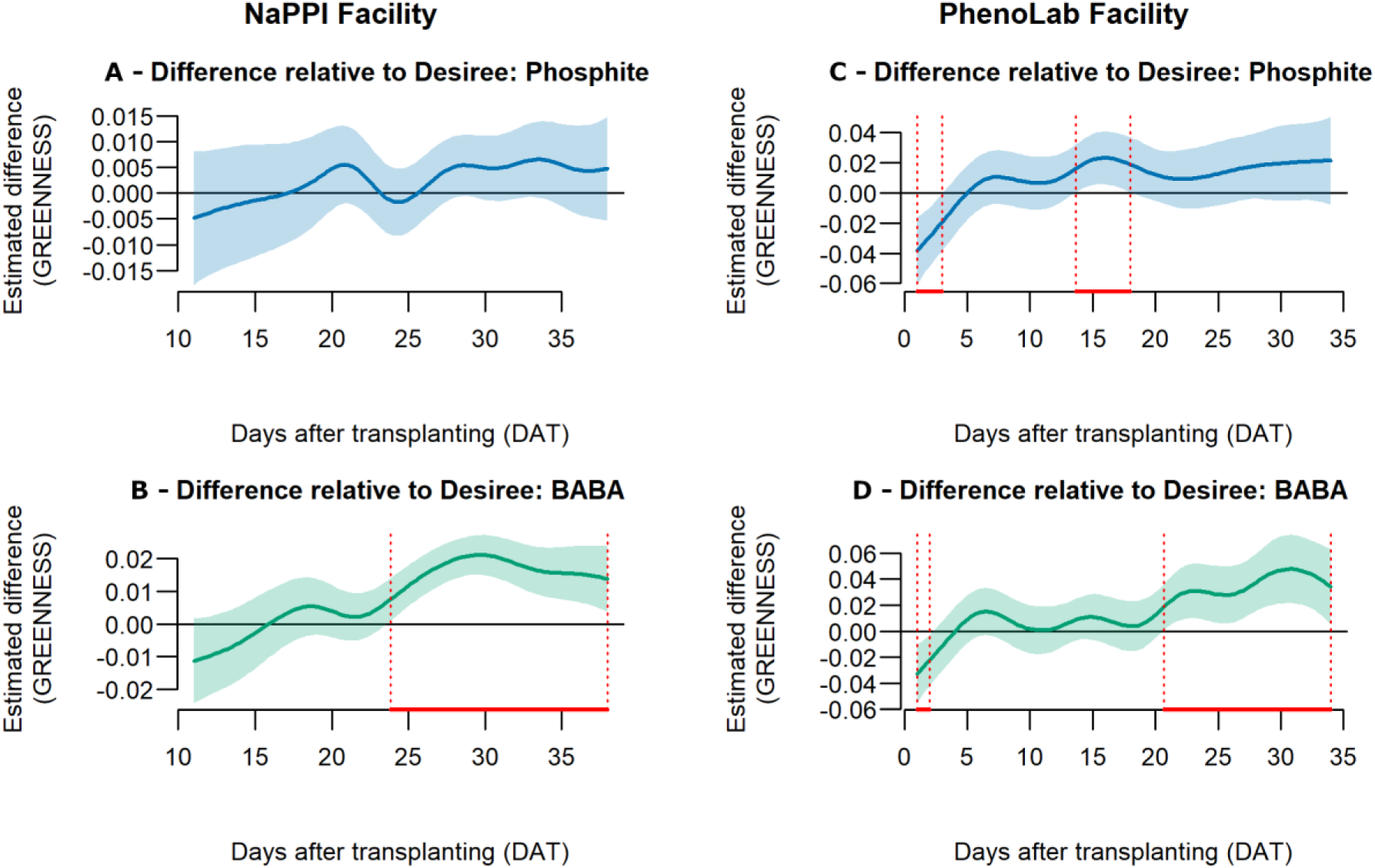
Greenness dynamics in response to induced-resistance treatments. Pointwise differences in greenness between the Desiree (cv.) control and phosphite-treated plants in NaPPI (A) or PhenoLab (C), and between the control and BABA-treated plants in NaPPI (B) or PhenoLab (D), are shown across days after transplanting (DAT). Group differences were evaluated using two-sided pairwise smooth-difference tests derived from the GAMMs, with significance concluded when the 95% confidence interval of the difference did not include zero. Six biological replicates per treatment were used, and IR treatments were applied weekly via foliar sprays. BABA-treated plants are shown in green, and phosphite-treated plants in blue. Red lines at the bottom of panels A–D indicate periods during which significant differences at the 95% confidence level were detected.

### Chlorophyll fluorescence measurements following IR treatments at the end of the experiment

To infer about general plant health, chlorophyll fluorescence measurements were conducted at the end of the experiment in NaPPI. Chlorophyll fluorescence measurements showed no significant differences between control and IR treatments for maximum PSII quantum yield (F_v_/F_m_), effective PSII quantum yield at light steady state (QY_Lss), or nonphotochemical quenching at light steady state (NPQ_Lss) (data not shown).

The fluorescence decline ratio (Rfd_Lss), used as an indicator of plant vitality, was significantly influenced by the treatments. BABA-treated plants exhibited lower Rfd_Lss values whereas KPhi-treated plants showed significantly higher values compared to the control (Fig. 4), indicating a contrasting physiological response between the two inducers.

**Fig. 4.**
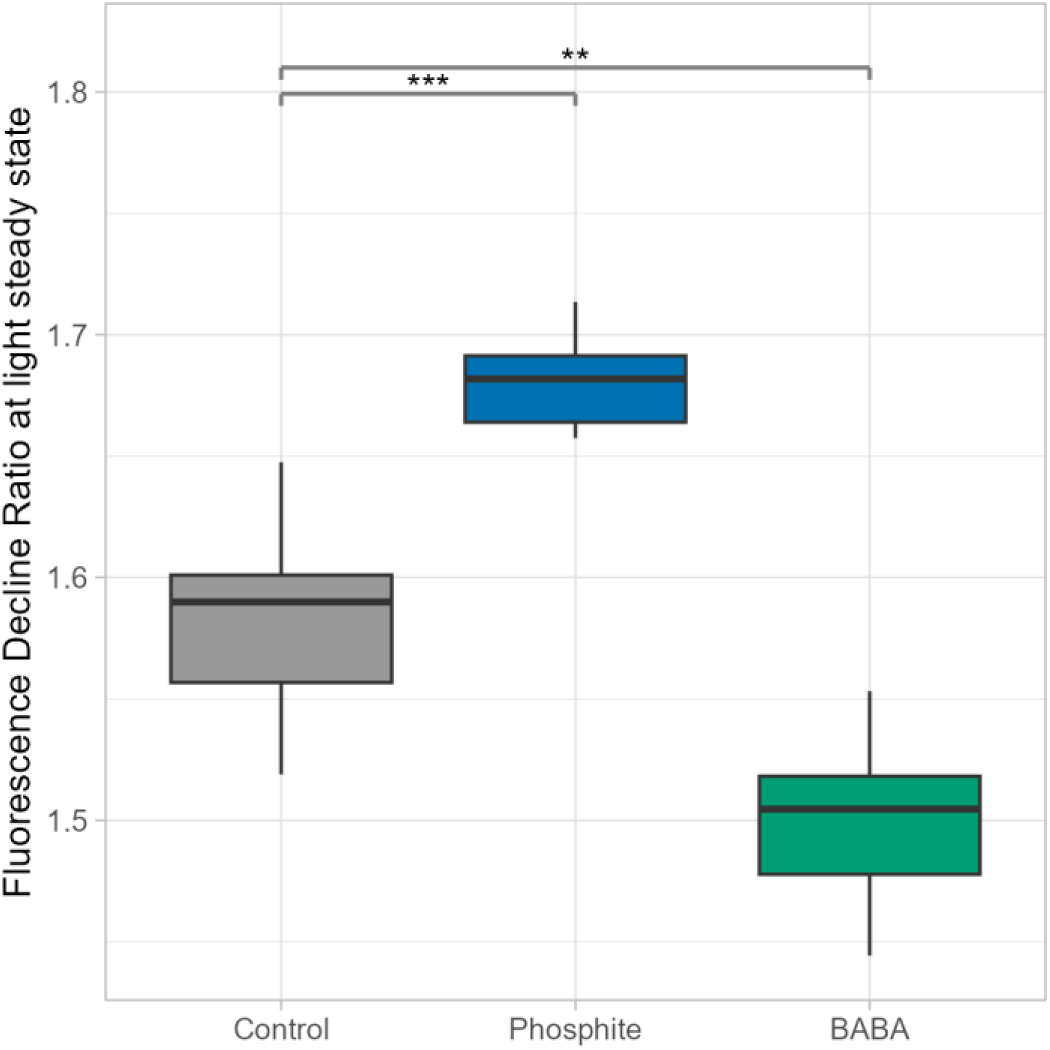
Rfd response to induced-resistance treatments at the end of the treatment. Boxplots show the distribution of the Fluorescence Decline Ratio at light steady state (Rfd_Lss) measured at 35–36 DAT in NaPPI. Six biological replicates per treatment were used. Group differences were assessed using two-sided Dunnett-type multiple-comparison tests applied to estimated marginal means (EMMs). Control plants are shown in grey, phosphite-treated plants in blue, and BABA-treated plants in green. Significant differences between IR treatments and the control are indicated by asterisks above each boxplot (*p < 0.05, **p < 0.01, ***p < 0.001).

### Continuous multispectral imaging following IR treatments

Multispectral imaging conducted in PhenoLab enabled calculation of the Normalized Difference Vegetation Index (NDVI), a spectral index commonly used as an indicator of canopy greenness and photosynthetic activity. Both BABA- and phosphite-treated plants maintained significantly higher NDVI values than the control during the early growth stage (Supplementary Fig. S1A). At later stages, NDVI values remained constant; while the control declined, BABA- and phosphite-treated plants remained relatively stable and slightly higher than the control, although these differences were not statistically significant (Supplementary Figs. S1B and S1C).

### Enzymatic analysis following IR treatments

Growth–defence trade-offs are assumed to occur in plants due to resource restrictions, which demand prioritization towards either growth or defence, respectively. Thus, we assessed the activity signatures of key enzymes of primary carbohydrate metabolism and antioxidant metabolism as related cell physiological proxy as key determinants of growth versus direct defence responses, respectively.

At the end of the experiment, the activities of eight antioxidant enzymes and twelve carbohydrate-metabolizing enzymes were quantified in both facilities (Table S1). Their contribution to treatment responses was evaluated using Principal Component Analysis (PCA). Treatments formed distinct clusters on the PCA plots in both facilities, indicating that enzymatic profiles were strongly associated with treatment effects (Fig. 5A). However, in NaPPI, variance in the control group was primarily associated with PK and AGPase, whereas in PhenoLab it was linked to CAT and PGM. For BABA-treated plants, NaPPI showed strong associations with PFK and PGM, while PhenoLab was linked to CWinv. In KPhi-treated plants, NaPPI was associated with HKX and CWinv, whereas PhenoLab correlated with PFK and Susy.

**Fig. 5.**
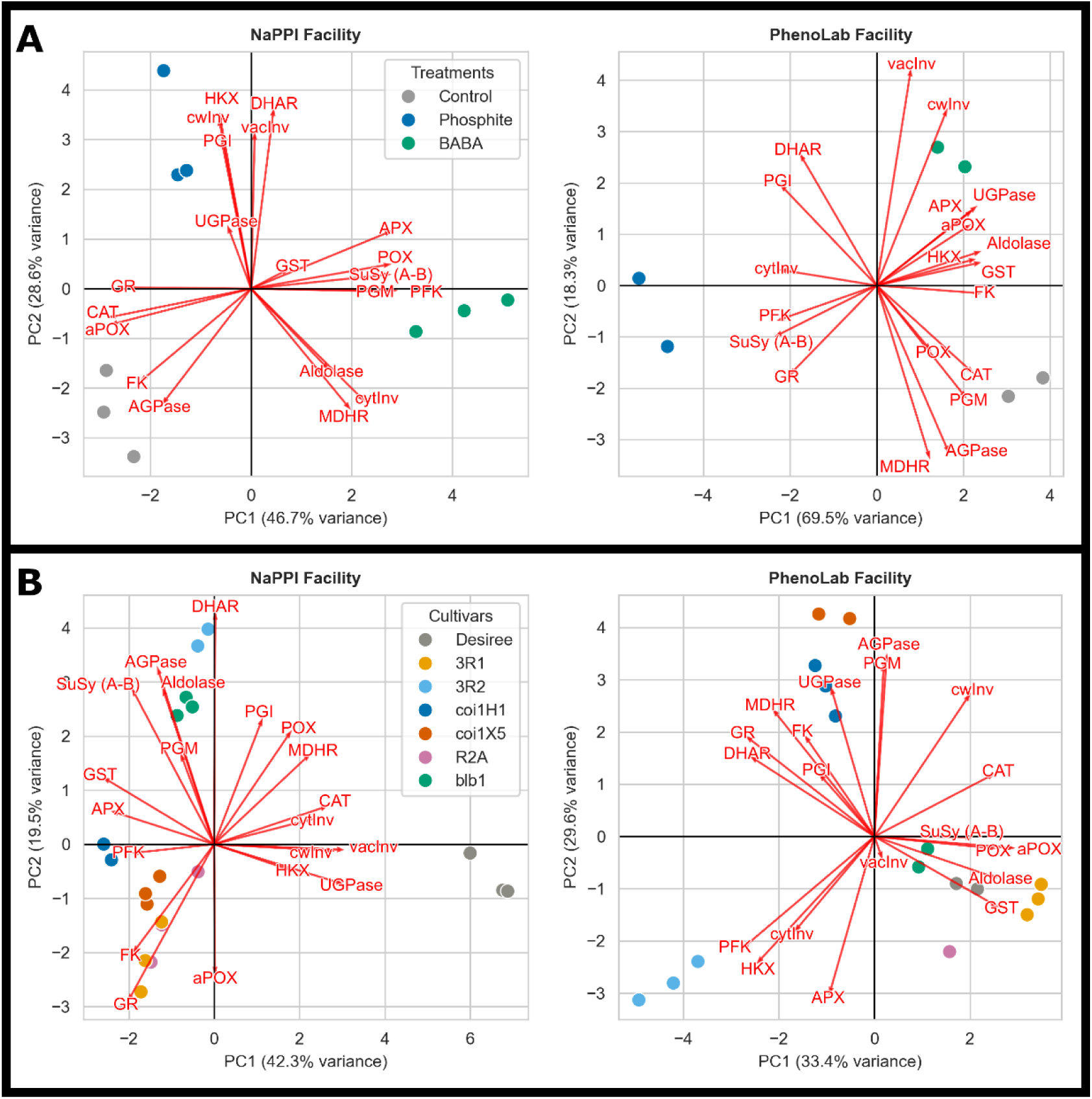
Principal Component Analysis (PCA) of enzymatic profiles in response to induced-resistance treatments and transgenic lines. Two-dimensional PCA scatter plots showing PC1 and PC2 based on endpoint enzymatic profiling from NaPPI (left panels) and PhenoLab (right panels). Induced-resistance treatments (A) include three biological replicates per treatment in NaPPI and two replicates in PhenoLab. Control plants are shown in grey, BABA-treated plants in blue, and phosphite-treated plants in green. Transgenic lines (B) include three biological replicates per line in both facilities. Control plants are shown in grey; transgenic lines are color-coded as follows: 3R1 (yellow), 3R2 (pale blue), coi1H1 (dark blue), coi1X5 (orange), R2A (purple), and blb1 (green). In all panels, red arrows indicate loading vectors representing the contribution and direction of each measured enzyme in the biplot.

In summary, the enzyme activity profiling revealed that in both facilities the IR treatments resulted in distinct signatures, supporting that those were affecting both primary carbohydrate metabolism and also antioxidant metabolism versus the mock controls. However, the key determinants of the differential responses were distinctly different in the two facilities. This was reflected in specific clustering when the data from both facilities were jointly analysed in a single PCA (Supplementary Fig. S2).

### End point measures of biomass following IR treatments

To determine if growth differences affected biomass or yield, final above-ground and tuber measurements were taken. BABA-treated plants in NaPPI showed a trend toward lower fresh weight than controls, a pattern not recorded in PhenoLab (Supplementary Fig. S3, upper panels). In contrast, dry weight in NaPPI was significantly reduced in these plants, while PhenoLab exhibited a similar but non-significant trend (Fig. 6). This pattern was opposite to the results for plant height but aligned with those for canopy area. No differences from the control were observed for KPhi-treated plants in either facility for both fresh and dry weight (Figs. S3 and 6), which was consistent with other growth measurements. Differences in plant growth did not translate into significantly changes of number of tubers or tuber weight compared to the control in either facility (Supplementary Fig. S3). Nonetheless, a common pattern was observed in both facilities, with tuber number tending to be higher in KPhi-treated plants in NaPPI and PhenoLab (Supplementary Fig. S3).

**Fig. 6.**
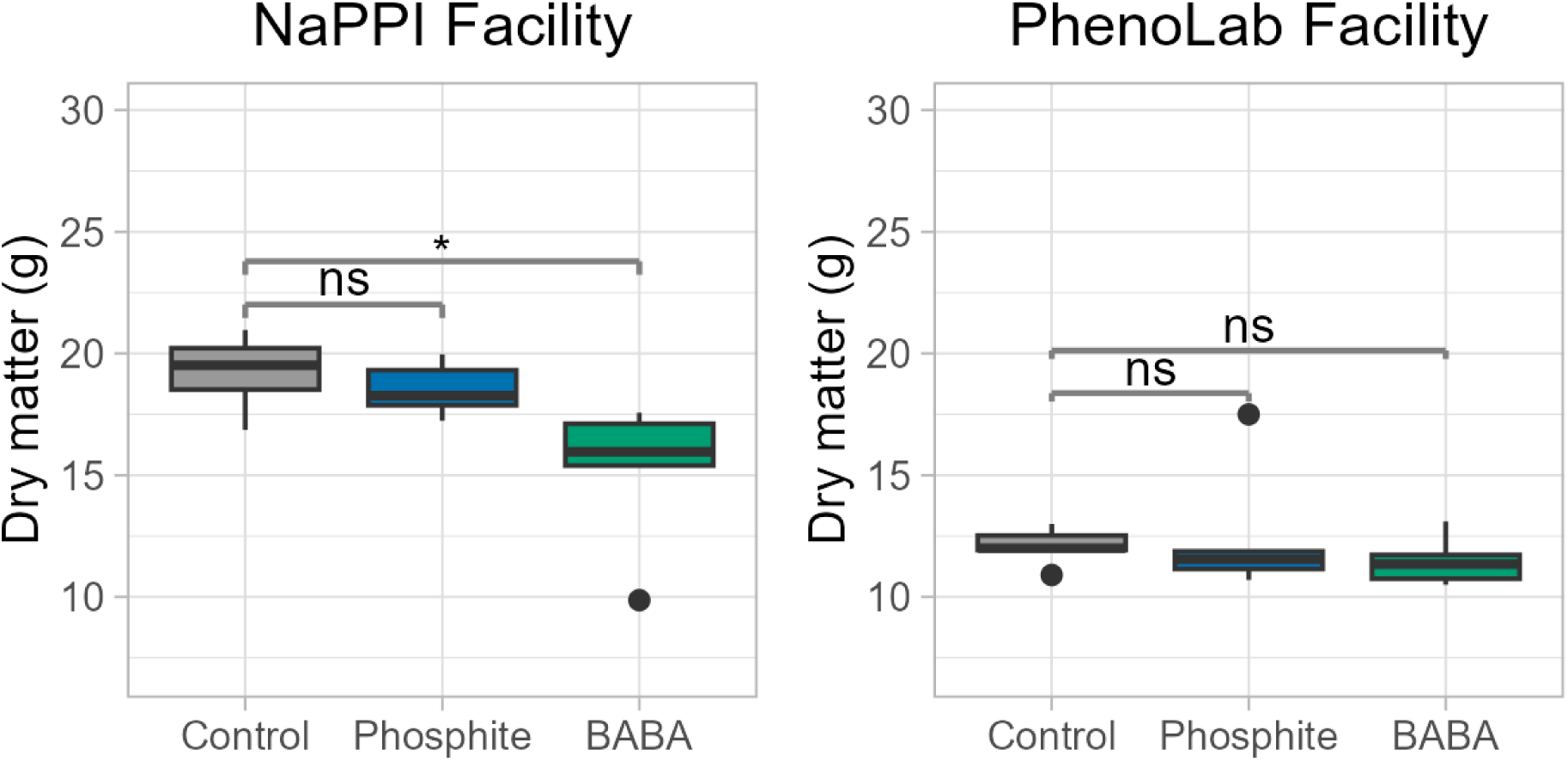
Dry matter response to induced-resistance treatments. Boxplots showing above-ground dry matter in NaPPI (left panel) and PhenoLab (right panel) at the end of the experiment. Each genotype includes six biological replicates. Group differences were assessed using two-sided Dunnett-type multiple-comparison tests applied to estimated marginal means (EMMs). Significant differences between IR treatments and the control are indicated by asterisks: *p < 0.05, **p < 0.01, ***p < 0.001.

### Transgenic lines with different resistance genes

In the second part of this study, we examined potential costs associated with resistance to *P. infestans* by evaluating four transgenic lines (cv. Désirée) - each carrying stacked Rpi genes or altered phytohormone sensing - across the two HTPP facilities over four weeks (Table 1). The broad-spectrum Rpi-blb1 gene from *S. bulbocastanum* provides resistance to multiple *P. infestans* races **(Van Der Vossen *et al*., 2003; Abreha *et al*., 2015)**, whereas line R2A carries the R2 gene, which recognises isolates expressing the Avr2 allele and therefore offers race-specific resistance **(Lenman *et al*., 2016)**. Line 3R2 contains three Rpi genes - two alleles from *S. bulbocastanum* (Rpi-blb1/RB and Rpi-blb2) and the broad-spectrum Rpi-vnt1.1 from *Solanum venturi* **(Ghislain *et al*., 2019) -** allowing assessment of potential additive or synergistic resistance effects under controlled physiological and developmental measurements.

### Plant height in transgenic lines over the course of the experiment

To assess treatment effects on plant height relative to the control, pointwise estimated differences were used. Lines expressing a single Rpi gene R2A showed no clear difference compared to the control in either facility (Supplementary Figs. S4A and C, top panels), whereas lines expressing the single Rpi gene blb1 exhibited intermittent periods of significantly lower plant height in NaPPI and sustained lower height from 16 to 34 DAT in PhenoLab (Supplementary Figs. S4B and D).

In NaPPI, lines with stacked Rpi genes showed contrasting patterns: 3R1 displayed no clear difference compared to the control, while 3R2 was significantly taller than the control from 34 to 38 DAT (Fig. 7A, top panels). Both the coi1 lines H1 and X5, which have reduced sensitivity to jasmonic acid methyl ester, were taller at the end of the experiment, with significant differences from 34 to 38 DAT (Fig. 7A, bottom panels). The same patterns were observed in PhenoLab, confirming these observations (Supplementary Fig. S5).

**Fig. 7.**
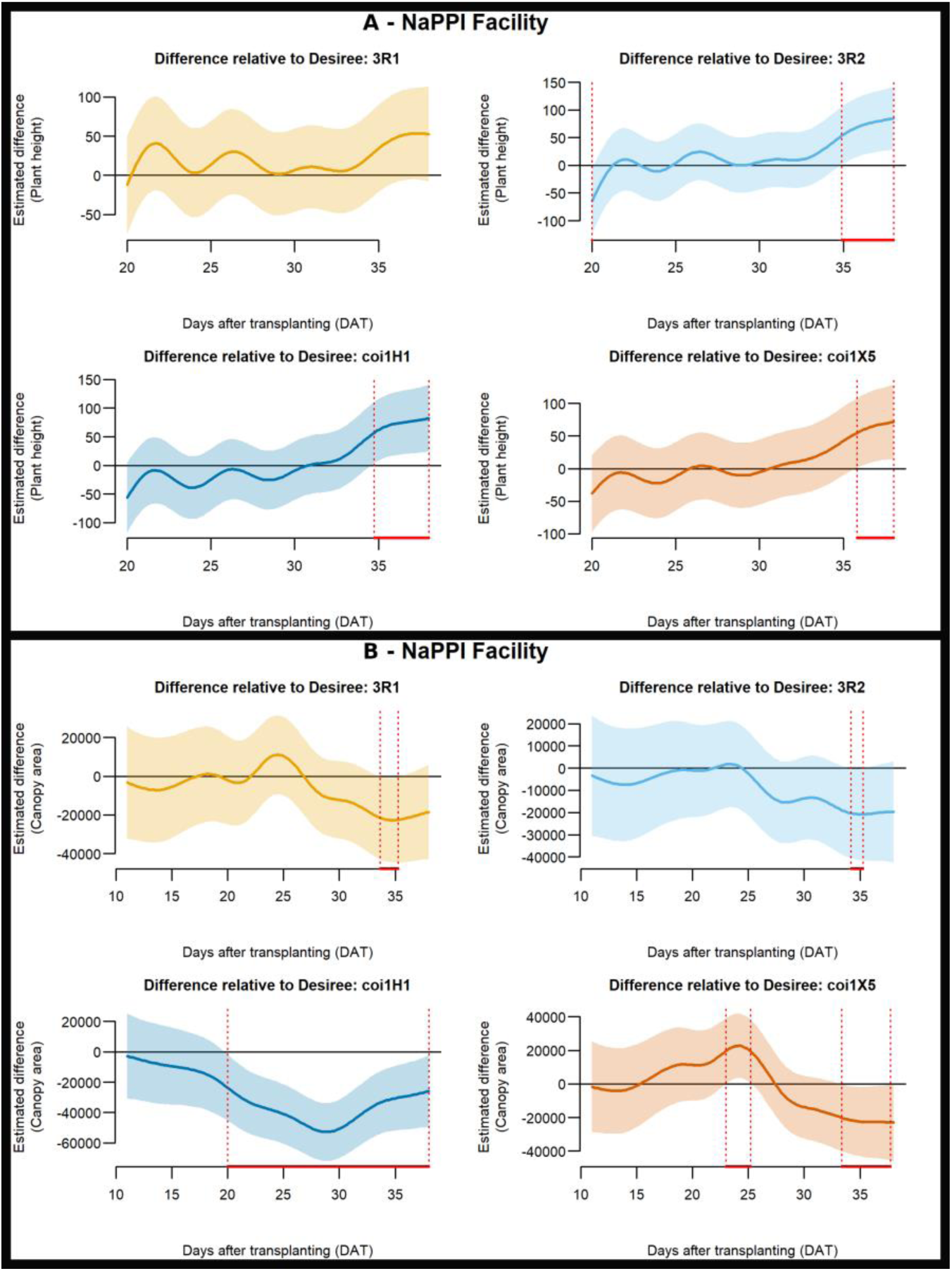
Plant height and canopy area dynamics in NaPPI for transgenic lines containing stacked Rpi genes (3R1, 3R2) or modified phytohormone pathways (coi1H1, coi1X5). Pointwise differences in plant height (A) and canopy area (B) between Desiree (cv.) and the transgenic lines 3R1 (yellow), 3R2 (pale blue), coi1H1 (dark blue), and coi1X5 (orange) are shown across days after transplanting (DAT). Group differences were evaluated using two-sided pairwise smooth-difference tests derived from the generalized additive mixed models (GAMMs), with significance concluded when the 95% confidence interval of the difference did not include zero. Six biological replicates per genotype were used. Red lines at the bottom of the plots indicate periods during which significant differences at the 95% confidence level were detected.

### Extracted plant canopy traits in transgenic lines over the course of the experiment

Total canopy area was measured for transgenic lines in NaPPI (Fig. 7B). Lines expressing single Rpi genes, R2A or blb1, showed no clear difference compared to the control (Supplementary Fig. S6). The coi1H1 line exhibited a significant lower canopy compared to control from 20 DAT until end of the growth period (Fig. 7B, bottom left panel), similar to BABA-treated Desiree (cv.). The coi1X5 line, however, showed a short significant higher canopy during the mid-course (from 23 to 25 DAT), followed by a significant lower canopy toward the end of the experiment (from 33 to 37 DAT; Fig. 7B, bottom right panel), a pattern resembling that of KPhi-treated plants. Other lines with stacked Rpi genes, 3R1 and 3R2, displayed similar trends to coi1X5 or KPhi-treated plants, with canopy area higher than the control during the mid-course and significantly lower toward the end of the experiment (from 33 to 35 DAT for 3R1 and 34 to 35 DAT for 3R2; Fig. 7B, upper panels).

### Color composition and chlorophyll fluorescence measurements in transgenic lines

The greenness index measured in transgenic lines in both facilities showed no significant differences compared to the control (data not shown). Similarly, chlorophyll fluorescence parameters measured in NaPPI, F_v_/F_m_, QY_Lss, and NPQ_Lss, did not differ significantly from control (data not shown). The plant vitality indicator Rfd_Lss in the coi1 mutant lines (H1 and X5) was comparable to the control (Supplementary Fig. S7). In contrast, Rfd_Lss tended to be slightly lower in lines carrying stacked Rpi genes (3R1 and 3R2) as well as in lines containing a single Rpi gene (R2A and blb1); however, these differences were not statistically significant (Supplementary Fig. S7)

### Continuous multispectral imaging of transgenic lines

NDVI measurements in PhenoLab showed that all transgenic lines increased until reaching a plateau (data not shown). Pointwise comparisons between each transgenic line and the control revealed that lines carrying stacked Rpi genes (3R1, 3R2) and those with modified phytohormone pathways (coi1H1, coi1X5) exhibited significantly higher NDVI values than the control during the early growth stage, similar to IR-treated plants (Supplementary Fig. S8). In contrast, lines expressing single Rpi genes (R2A and blb1) did not display such early NDVI differences compared to the control (Supplementary Fig. S8).

### Enzymatic analysis in transgenic lines

Principal Component Analysis (PCA) was performed to evaluate the contribution of enzyme activities to variation among transgenic lines across the two facilities (Fig. 5B). In NaPPI, control samples clustered opposite to most transgenic lines, primarily associated with high loadings of Invertase and UGPase. Lines carrying stacked Rpi genes (3R1, 3R2) segregated in distinct directions: variance in 3R1 was mainly explained by PK and GR, whereas 3R2 was associated with DHAR. Transgenic lines with modified phytohormone pathways (coi1H1, coi1X5) clustered closely together, strongly correlated with PFK. Single Rpi gene lines (R2A, blb1) showed contrasting patterns: R2A grouped near 3R1, while blb1 clustered with 3R2 (Fig. 5B, left panel).

In PhenoLab, clustering patterns differed markedly. Control samples grouped together with blb1 and 3R1, associated with Aldolase and GST. The other stacked Rpi line (3R2) formed a distinct cluster linked to PFK and HKX, while R2A also separated independently. Both coi lines (coi1H1, coi1X5) again clustered together, associated with UGPase and AGPase (Fig. 5B, right panel).

In summary, the enzyme activity profile revealed that in both facilities the different transgene mediated resistance resulted in distinct biosignature, supporting that those were differentially affecting both primary carbohydrate metabolism and also antioxidant metabolism in relation to the mock control. However, the key determinants of the differential responses were completely different in the two facilities. This was reflected in a distinct, facility specific clustering when the data from both facilities were jointly analysed in a single PCA (Supplemental Fig. S2).

### End point measures of biomass in transgenic lines

Across both facilities, patterns of biomass and yield responses varied by genotype and environment (Figs. 8 and S9). In NaPPI, most transgenic lines exhibited lower fresh and dry biomass compared to the control, although these differences were generally not statistically significant, except for coi1X5, which showed a significantly lower dry weight (Fig. 8 left panel). Regarding yield, 3R1 produced a significantly greater number of tubers, and coi1X5 had a significantly higher tuber weight, while all other lines showed no significant differences in tuber number or weight compared to the control (Supplementary Fig. S9 middle and bottom panels of the left side). In PhenoLab, significant reductions in biomass were observed for lines carrying stacked Rpi genes (3R1, 3R2) and single Rpi genes (R2A, blb1), with fresh weight significantly reduced for 3R1, 3R2, and blb1, and dry weight significantly reduced for all four lines. On the contrary, coi1H1 displayed a significant increase in both fresh and dry weight, while coi1X5 showed a non-significant trend toward increased dry weight (Fig. 8 right panel). Tuber number tended to be higher across transgenic lines, but only 3R2 and R2A were significantly greater than the control. Tuber weight varied among lines, higher in 3R1 and lower in coi1H1, blb1, 3R2, coi1X5, and R2A, but none of these differences were statistically significant (Supplementary Fig. S9, bottom right panel). Overall, these results suggest that, depending on the trait measured, several transgenic lines showed indications of reduced above-ground biomass compared to the Désirée wild type, pointing to potential growth costs associated with resistance gene expression. However, for most lines, tuber number and weight did not differ significantly from the control, with the notable exception of coi1X5, which produced higher tuber biomass in NaPPI.

**Fig. 8.**
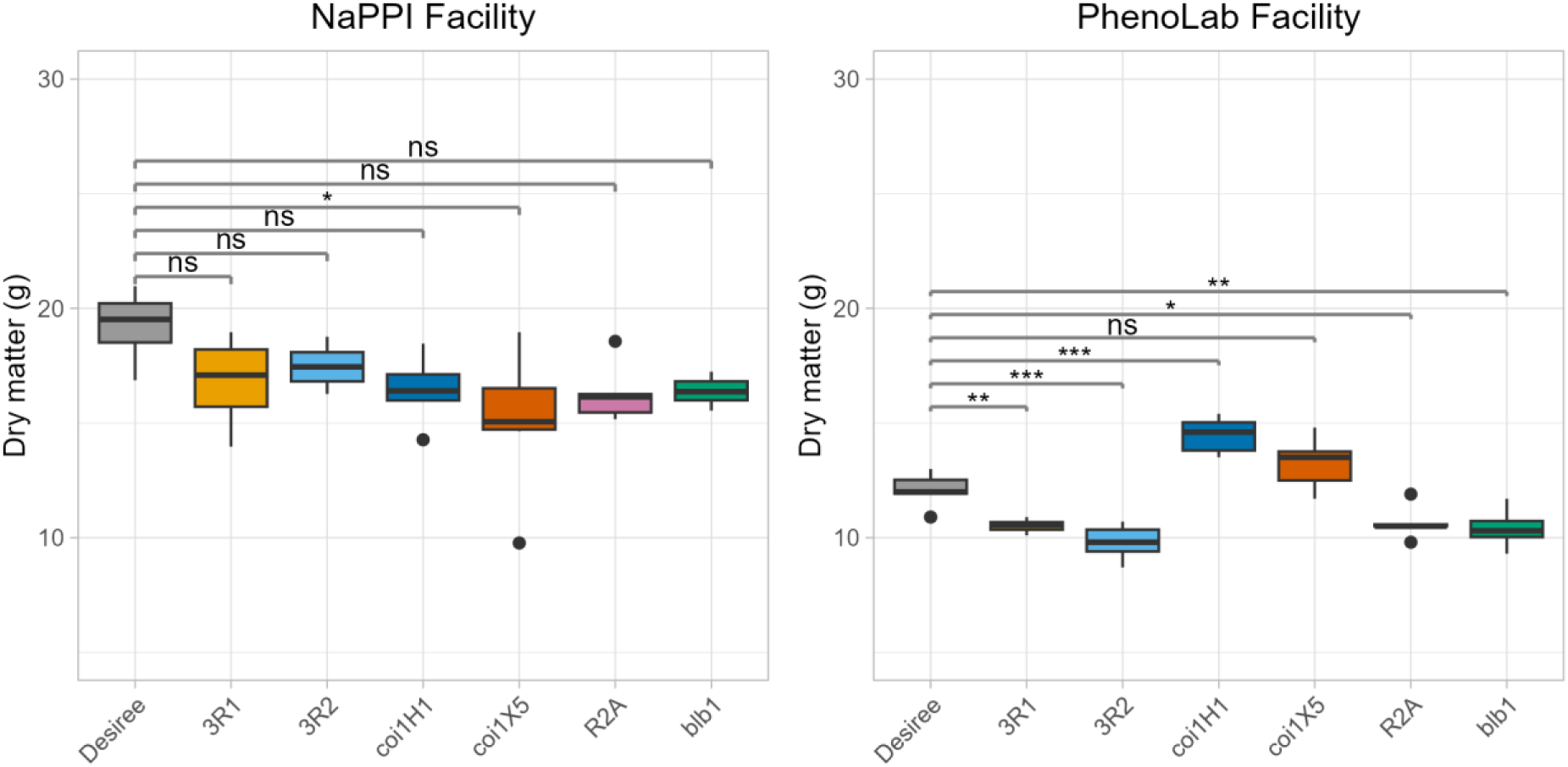
Dry matter response in transgenic lines containing stacked Rpi genes (3R1, 3R2), modified phytohormone pathways (coi1H1, coi1X5), and single Rpi genes (R2A, blb1). Boxplots showing above-ground dry matter in NaPPI (left panel) and PhenoLab (right panel) at the end of the experiment. Each genotype includes six biological replicates. Group differences were assessed using two-sided Dunnett-type multiple-comparison tests applied to estimated marginal means (EMMs). Significant differences between IR treatments and the control are indicated by asterisks: *p < 0.05, **p < 0.01, ***p < 0.001.

## Discussion

Recent advances on sensor-based imaging and automation technology allows high-throughput phenotyping for multiple traits related to plant development under different stress conditions. It is less used to assess fitness costs as a consequence of different defence strategies or for monitoring the effects of biostimulants. In this study, we used image-based sensors to assess plant growth, and morphological and physiological changes over time in transgenic potato lines overexpressing various resistance genes to *P. infestans* or with reduced sensitivity to phytohormones, and in response to induced resistance (IR) triggered by BABA and potassium phosphite (KPhi) in the absence of the pathogen. We found morphological and physiological changes in both the transgenic lines and in response to IR treatment suggesting that these phenotyping methods may be useful for investigating potential costs triggered by the presence of resistance genes or IR.

There are few studies where the morphology of resistance genes including Rpi-blb1 has been studied over time in such detail. We have previously used these specific potato transgenic Rpi-blb1 lines and did not notice a clear morphological difference **(Abreha *et al*., 2015)**. The 3R gene construct was evaluated during the growth season and no statistical difference in morphology was detected between the transgene and its genetic background by weekly measurements in the field **(Byarugaba *et al*., 2021)**, suggesting that environmental conditions outweigh genotypic effects on plant morphology.

The coi1 gene was knocked down by RNAi in lines H1 and X5, making them less sensitive to jasmonic acid methyl ester and enabling us to assess allocation costs when hormone signalling is disrupted. While specific studies detailing the effects of the coi transgene on potato height are lacking, the literature indicates that transgene expressions, particularly those that have the potential to influence hormonal pathways or metabolic processes, can result in dwarfing effects or altered height. In *Arabidopsis thaliana* and *Nicotiana benthamiana*, **Noir et al. (2013)** observed that MeJA inhibits leaf growth through its receptor, COI1, by reducing the number and size of cells, which ultimately affects overall plant height. Further supporting the effect of jasmonates, **Ravnikar et al. (1992)** reported that jasmonic acid treatments influenced shoot and root development in *Solanum tuberosum*, resulting in increased growth under specific concentrations of jasmonic acid. Their findings emphasize the potential growth-promoting effects of jasmonates. While most of this specific research focuses on species outside the Solanum genus, it suggests that similar mechanisms may operate in the Solanum genus, indicating broader implications of jasmonate signaling.

To activate disease resistance in *Arabidopsis*, the biologically active R-enantiomer of BABA binds to the aspartyl tRNA synthetase (AspRS) IBI1 resulting in the accumulation of uncharged tRNAAsp and, consequently, plant stress and growth reduction occurs if BABA is applied in high concentrations **(Luna *et al*., 2014)**. We used a BABA concentration of 10 mM for being previously shown to be an effective concentration against potato late blight and it has previously confirmed that at this concentration there were changes in transcript expression and protein secretion in potato plants **(Bengtsson *et al*., 2014*a*,*b*)**. After four weekly BABA treatments, we observed reduced biomass growth in potato plants, with decreases in both top view area and perimeter. The results go along with observations in *Arabidopsis thaliana* where BABA-IR shifted to a BABA-induced stress status which was hypothesized to be linked to the concentrations used **(Van Hulten *et al*., 2006)**. At the same time, we found that treating potato plants with BABA boosted the plant height.

There is evidence that phosphite applications can influence plant growth and architecture. The impact of phosphite on various aspects of plant development, including morphology, photosynthesis, and resistance to pathogens, has been documented in several studies **(Mohammed *et al*., 2022; Pérez-Zavala *et al*., 2024)**. Furthermore, we previously showed that phosphite BABA treatment of potato cv. Désirée at the same amounts as the present study led to a 40% overlap of differentially expressed genes of KPhi and BABA treatment **(Burra *et al*., 2014)**. Nevertheless, the repeated treatment of phosphite did not provide morphological significant differences compared to untreated control. The lack of negative effects on plant growth and development highlights KPhi as a suitable plant protection agent, as it can be applied at levels giving protection against *P. infestans* without compromising plant performance.

Chlorophyll fluorescence is commonly affected when plants are under stress **(Kalaji *et al*., 2016; Pérez-Bueno *et al*., 2019).** In this study, BABA and phosphite treatments did not significantly alter the maximum quantum efficiency of PSII photochemistry (F_v_/F_m_), a robust photosynthetic efficiency indicator. Small changes on F_v_/F_m_ values were observed over time, but always within limits of those from a healthy plant tissue. Since measuring photosynthetic rate in the field is time-consuming, studies have identified a strong correlation between leaf greenness and photosynthetic rate **(Signorelli *et al*., 2025)**.

The observed phenotypic effects can be interpreted within the broader growth–defence trade-off framework, where defence activation imposes allocation costs through pleiotropic gene effects and antagonistic hormonal cross-talk, particularly between salicylate- and jasmonate-mediated pathways **(Brown and Rant, 2013; Karasov *et al*., 2017; He *et al*., 2022)**. Under pathogen-free conditions, BABA treatment produced the most pronounced phenotypic signatures, including reduced canopy area, lower dry biomass and altered chlorophyll fluorescence, consistent with a measurable but modest allocation cost. In contrast, KPhi and most transgenic Rpi lines showed minimal growth penalties, suggesting that not all defence mechanisms impose equivalent trade-offs, an observation aligned with the notion that fitness costs depend on the specific signalling pathway engaged and its degree of constitutive activation. The subtlety of these trade-offs is also consistent with the argument that crop breeding may have partially buffered the growth–defence conflict relative to model plants such as Arabidopsis **(Van Hulten *et al*., 2006)**, and with recent evidence that R gene expression costs arise mainly from downstream signalling rather than from gene expression itself **(von Dahlen *et al*., 2023).**

The comparative assessment of the impact of the IR treatments and expression of resistance inducing transgenes on key enzymes of carbohydrate an antioxidant metabolism revealed in the two facilities distinct impacts on these cells physiological proxy **(Großkinsky *et al*., 2015; Jammer *et al*., 2022)**. However, the signatures were completely different in the two facilities within greenhouses, indicating, that these signatures are modulated by the specific external factors such as illumination and external photoperiod. In fact, the experiments were conducted in May-June (NaPPI) and in November-December (PhenoLab) resulting in clear temperature differences (see below). Thus, no specific IR treatment or transgene specific signatures could be identified in the comparative analyses of experiments across the facilities in Denmark and Finland.

The use of two independent phenotyping platforms, NaPPI and PhenoLab, provides significant methodological value by enabling the systematic benchmarking of high-throughput systems and ensuring the reproducibility of results across different research environments. Our findings demonstrate a high level of robustness for the two shared phenotypic traits: plant height and greenness, which exhibited consistent treatment-specific trajectories in both Finland and Denmark despite variations in local environmental controls.

This consistency is noteworthy given the differences in environmental control and cultivation conditions between the two facilities. At NaPPI, plants were grown in 5 L pots under slightly warmer, drier conditions with marginally higher light intensity (day averages of 25.6 °C, 37.5% RH; 250 µmol m^-2^ s^-1^), whereas at PhenoLab they were maintained in 1.5 L pots under cooler, more humid conditions with slightly lower light (22.8 °C, 56.9% RH; 224.3 µmol m^-2^ s^-1^). In addition, NaPPI showed greater thermal variability, with daytime temperatures reaching 35.9 °C, while PhenoLab remained more stable, with a maximum of 25.1 °C. Despite these contrasting conditions, the significant increase in plant height and the elevated greenness index, reflecting a paler color, following BABA treatment was successfully validated across both facilities. Conversely, traits such as enzymatic activity and final biomass appeared more sensitive to facility-specific conditions. Principal Component Analysis (PCA) of enzymatic signatures revealed distinct clustering patterns for each site, suggesting that local environmental factors can modulate or even override specific metabolic responses to treatments.

In this study, we demonstrated that high-throughput phenotyping can provide new insights into the interaction between plants and resistance inducers, and that subtle differences in growth can be robustly detected. Notably, we employed two independent phenotyping platforms and directly compared results across both, a unique approach that strengthens the reliability and generalisability of our findings. Ultimately, this cross-facility approach is critical for distinguishing biologically meaningful signals from technical variability, thereby strengthening the reliability and generalisability of the detected trade-offs associated with potato defense mechanisms. Our analyses indicate that at optimum concentrations for protection against potato late blight, BABA and phosphite could be integrated into disease management strategies without substantial costs to potato plants. High-throughput phenotyping platforms thus represent a powerful tool for quantifying potential fitness costs associated with novel resistance genes, resistance mechanisms, or induced resistance strategies—including biological control agents—by enabling precise, non-destructive assessment of plant growth and developmental trade-offs across different experimental systems. However, whether these subtle changes translate to impacts on crop production and yield at larger scales remains to be determined and will require further testing under field conditions across diverse environments and multiple years.

## Supplementary data

**Table S1. Endpoint enzymatic activity data used for principal component analysis of induced-resistance treatments and transgenic lines across NaPPI and PhenoLab.**

**Fig. S1.**
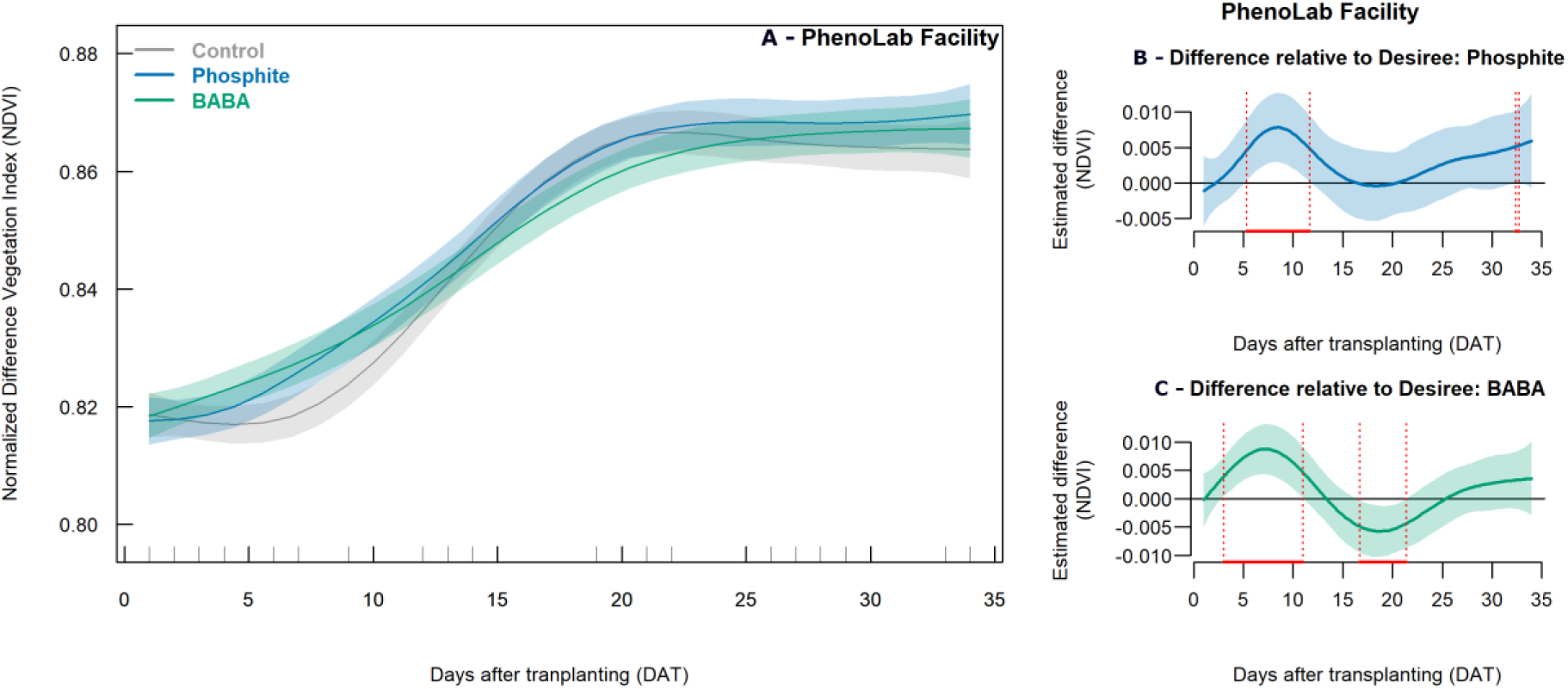
NDVI dynamics in response to induced-resistance treatments. Generalized Additive Mixed Models (GAMMs) were used to analyse NDVI measured in PhenoLab (A). Pointwise differences between the NDVI of the Desiree (cv.) control and phosphite-treated plants (B), or BABA-treated plants (C), are shown across days after transplanting (DAT). Group differences were evaluated using two-sided pairwise smooth-difference tests derived from the GAMMs, with significance concluded when the 95% confidence interval of the difference did not include zero. Six biological replicates per treatment were used, and IR treatments were applied weekly via foliar sprays. Control plants are shown in grey, BABA-treated plants in green, and phosphite-treated plants in blue. Shaded bands represent the 95% confidence intervals of the GAMM fits. Red lines at the bottom of panels B and C indicate periods during which significant differences at the 95% confidence level were detected.

**Fig. S2.**
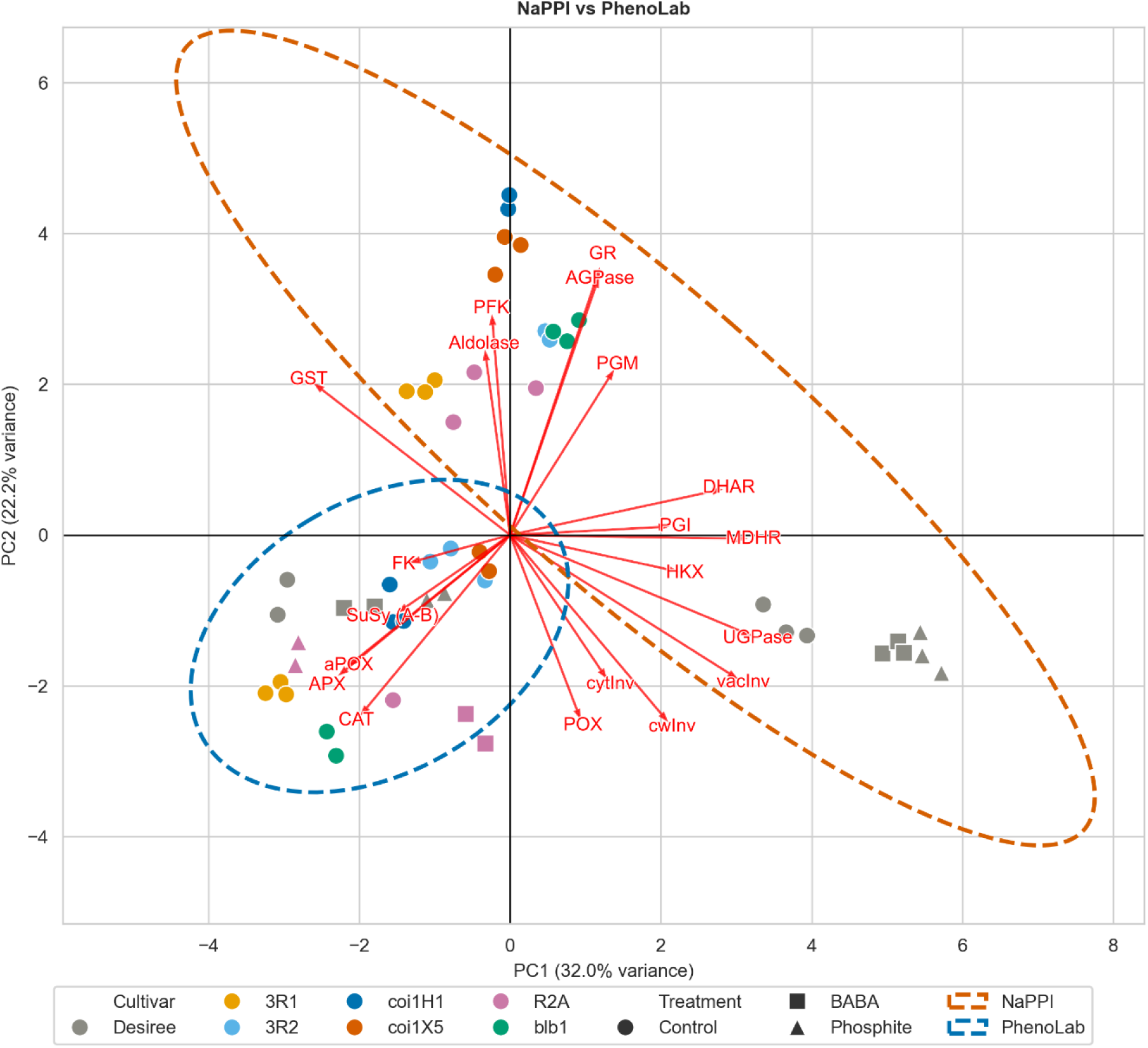
Principal Component Analysis (PCA) of enzymatic profiles combining induced-resistance treatments and transgenic lines across facilities. Two-dimensional PCA biplot showing PC1 and PC2 based on endpoint enzymatic profiling from NaPPI and PhenoLab combined into a single analysis. The biplot includes induced-resistance treatments (Control, BABA, phosphite) and transgenic lines (3R1, 3R2, coi1H1, coi1X5, R2A, blb1), together with the control cultivar Desiree. Control plants are shown in grey; transgenic lines are color-coded as follows: 3R1 (yellow), 3R2 (pale blue), coi1H1 (dark blue), coi1X5 (orange), R2A (purple), and blb1 (green) and shaped by treatment (circles: Control; squares: BABA; triangles: phosphite). Dashed ellipses indicate clustering by facility (NaPPI: orange dashed line; PhenoLab: blue dashed line). Red arrows represent loading vectors indicating the contribution and direction of each measured enzyme in the biplot.

**Fig. S3.**
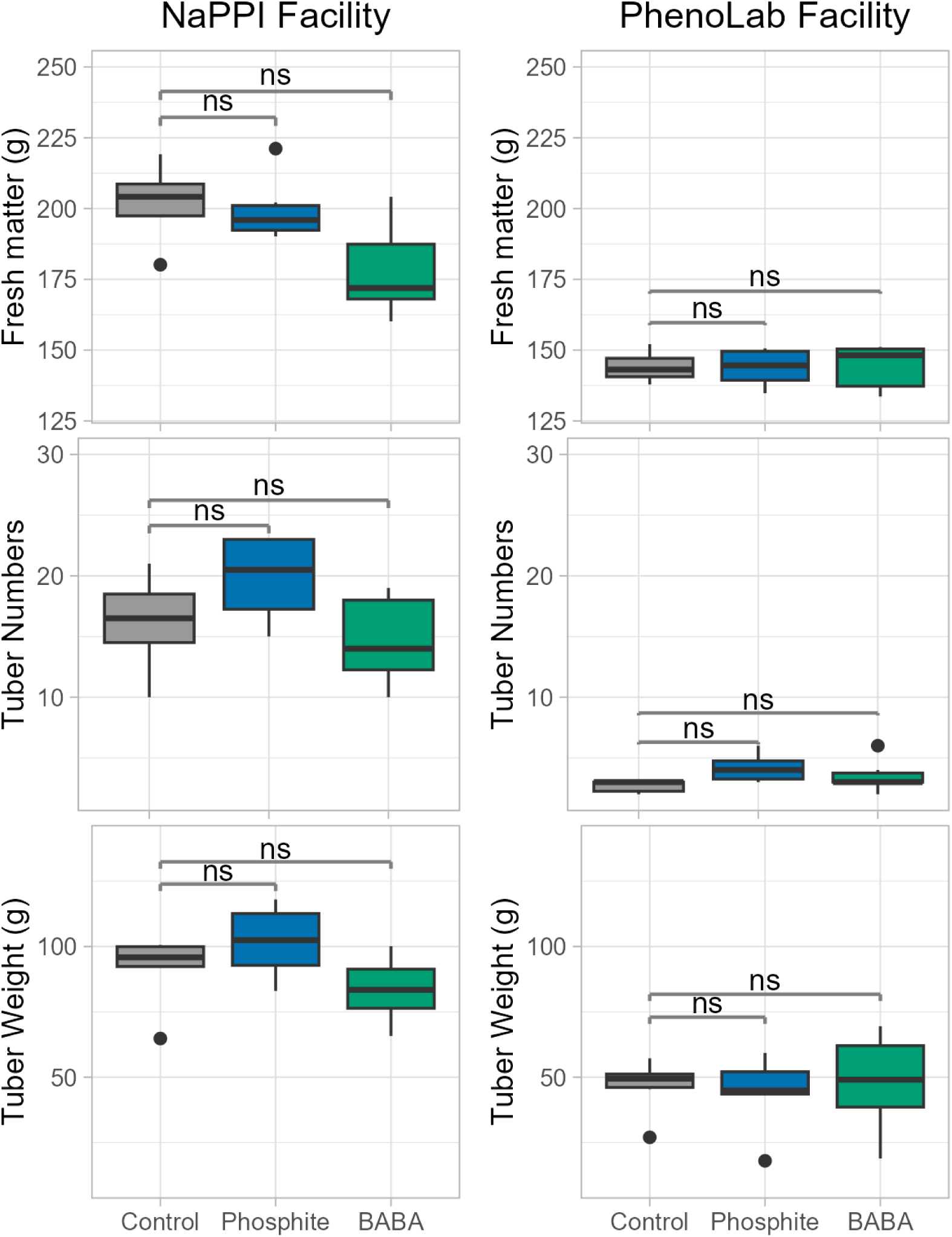
Yield parameters response to induced-resistance treatments. Boxplots showing above-ground fresh weight (top panels), number of tubers (middle panels), and total tuber weight (lower panels) for plants grown in NaPPI (left panels) and PhenoLab (right panels) at the end of the experiment. Each genotype includes six biological replicates. Group differences were assessed using two-sided Dunnett-type multiple-comparison tests applied to estimated marginal means (EMMs). Significant differences between IR treatments and the control are indicated by asterisks: *p < 0.05, **p < 0.01, ***p < 0.001.

**Fig. S4.**
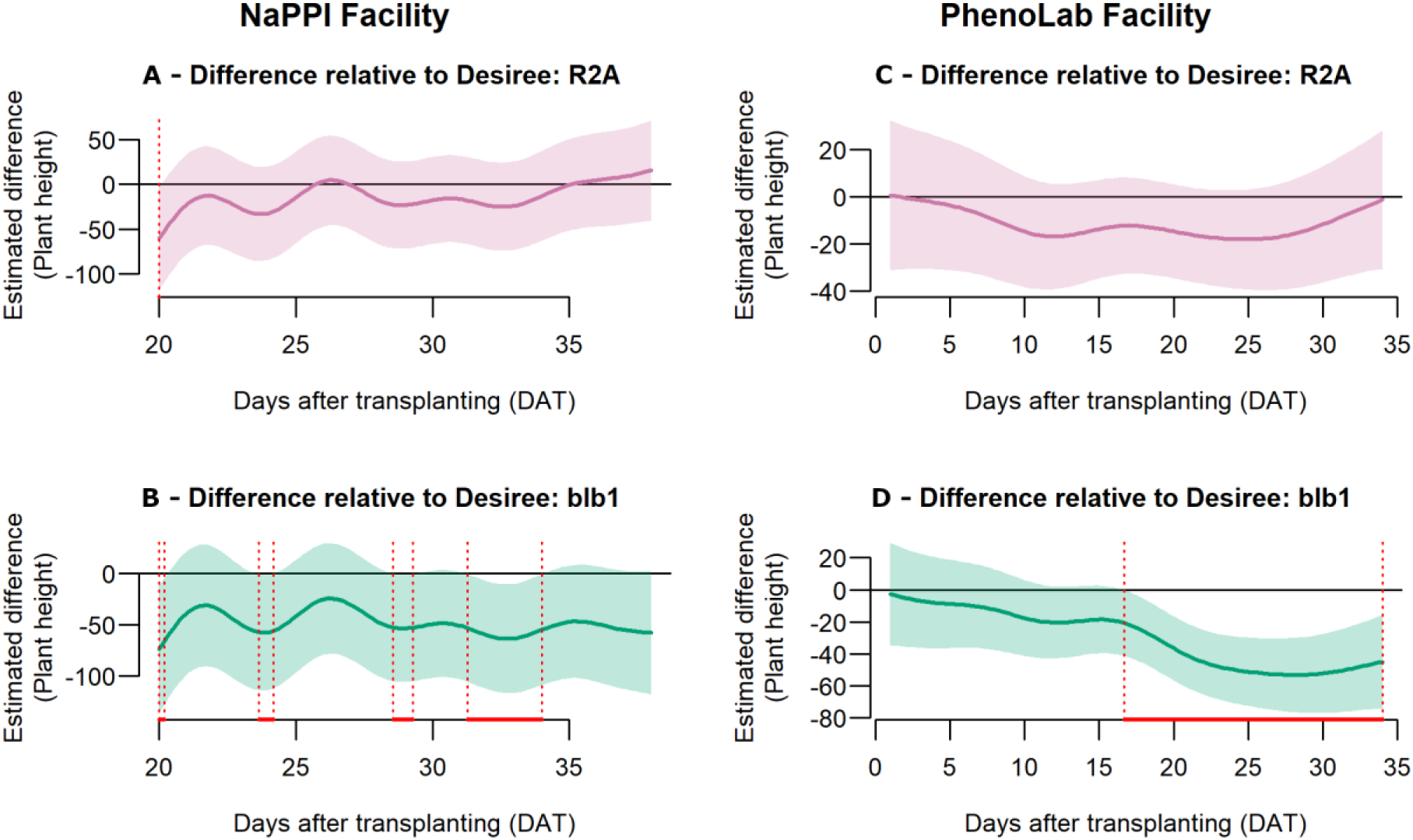
Plant height dynamics in transgenic lines expressing single Rpi genes (R2A, blb1). Pointwise differences in plant height between Desiree (cv.) and the transgenic lines R2A (purple) and blb1 (green) are shown for NaPPI (A–B) and PhenoLab (C–D) across days after transplanting (DAT). Group differences were evaluated using two-sided pairwise smooth-difference tests derived from the generalized additive mixed models (GAMMs), with significance concluded when the 95% confidence interval of the difference did not include zero. Six biological replicates per genotype were used. Red lines at the bottom of each plot indicate periods during which significant differences at the 95% confidence level were detected.

**Fig. S5.**
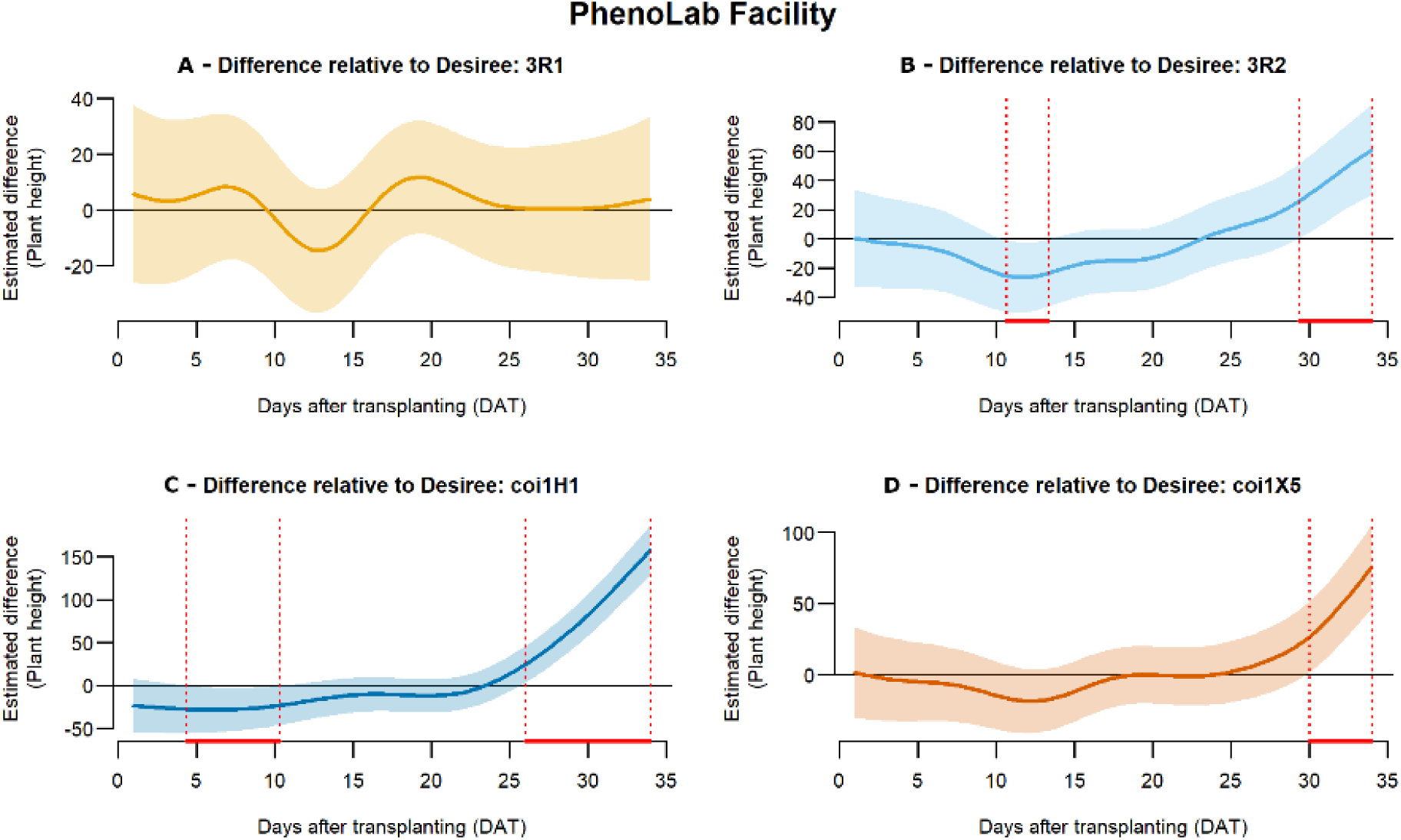
Plant height dynamics in Phenolab for transgenic lines containing stacked Rpi genes (3R1, 3R2) or modified phytohormone pathways (coi1H1, coi1X5). Pointwise differences in plant height between Desiree (cv.) and the transgenic lines 3R1 (A, yellow), 3R2 (B, pale blue), coi1H1 (C, dark blue), and coi1X5 (D, orange) are shown across days after transplanting (DAT). Group differences were evaluated using two-sided pairwise smooth-difference tests derived from the generalized additive mixed models (GAMMs), with significance concluded when the 95% confidence interval of the difference did not include zero. Six biological replicates per genotype were used. Red lines at the bottom of the plots indicate periods during which significant differences at the 95% confidence level were detected.

**Fig. S6.**
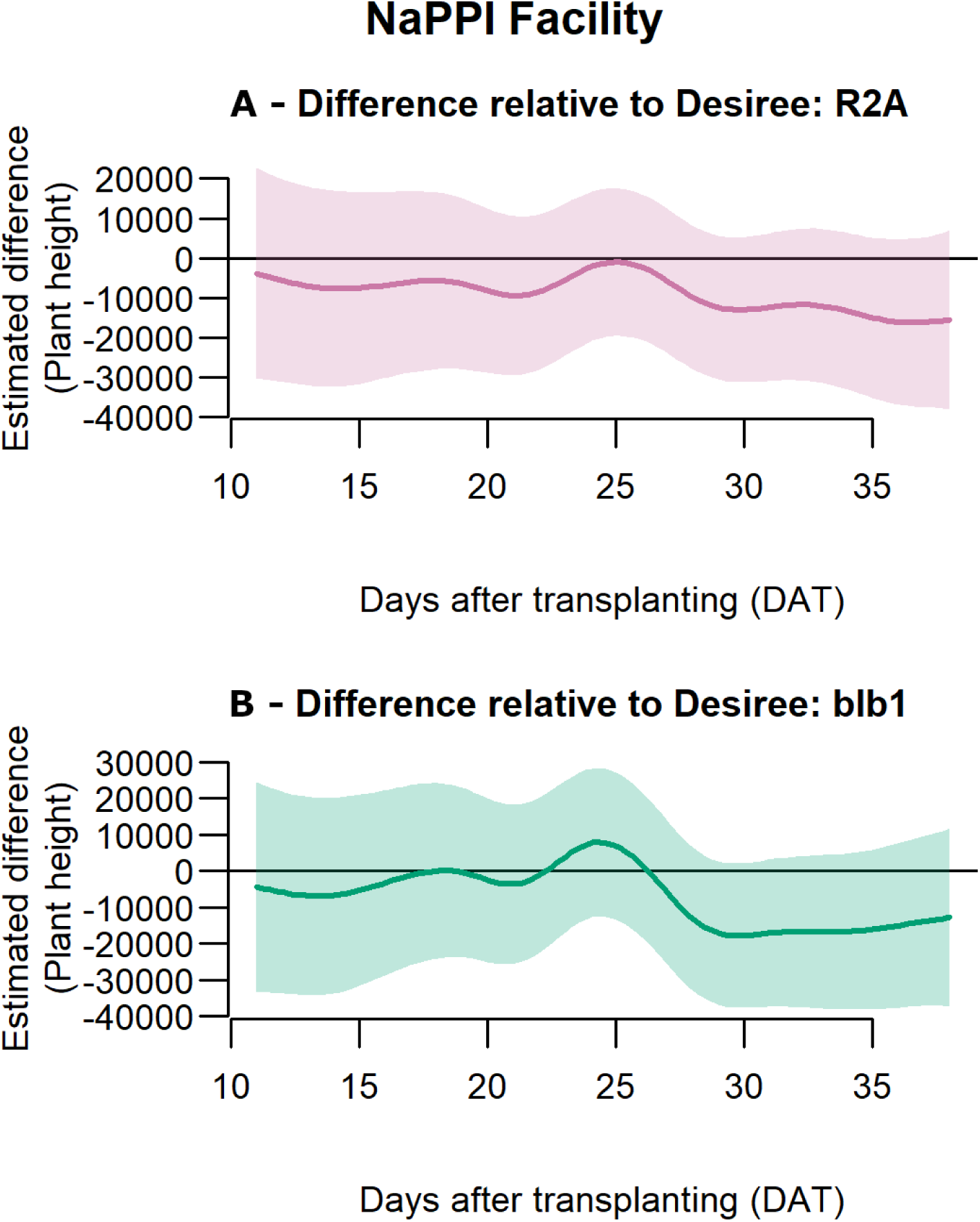
Plant height and canopy area dynamics in NaPPI for transgenic lines expressing single Rpi genes (R2A, blb1). Pointwise differences in canopy area between Desiree (cv.) and the transgenic lines R2A (A, purple) and blb1 (B, green) are shown across days after transplanting (DAT) in NaPPI. Group differences were evaluated using two-sided pairwise smooth-difference tests derived from the generalized additive mixed models (GAMMs), with significance concluded when the 95% confidence interval of the difference did not include zero. Six biological replicates per genotype were used. Red lines at the bottom of the plots indicate periods during which significant differences at the 95% confidence level were detected.

**Fig. S7.**
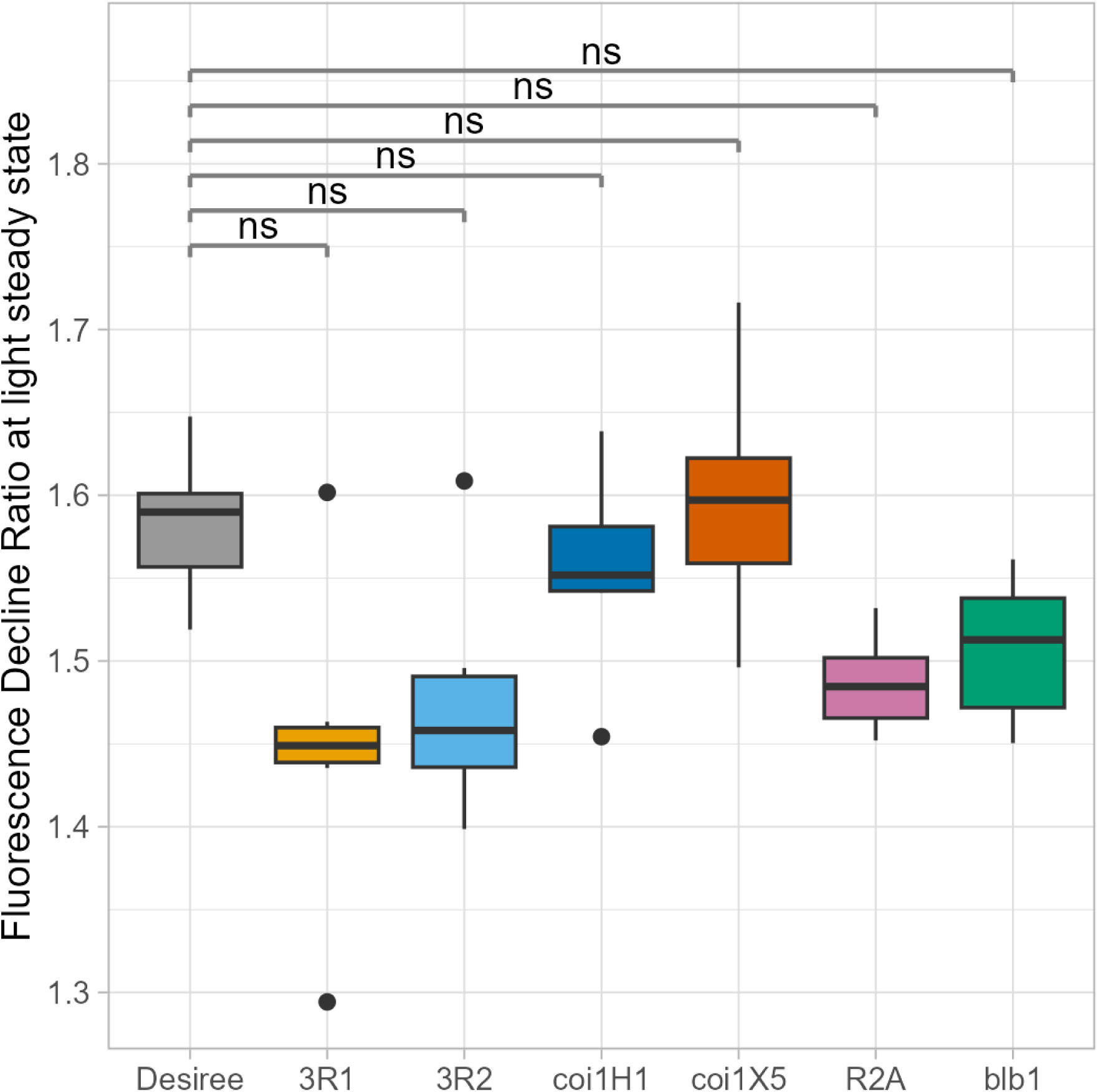
Rfd response in NaPPI for transgenic genic lines containing stacked Rpi genes (3R1, 3R2), modified phytohormone pathways (coi1H1, coi1X5), and single Rpi genes (R2A, blb1). Boxplots show the Fluorescence Decline Ratio at light steady state (Rfd_Lss) measured at 35–36 DAT in NaPPI. Six biological replicates per treatment were used. Control plants are shown in grey; transgenic lines are color-coded as follows: 3R1 (yellow), 3R2 (pale blue), coi1H1 (dark blue), coi1X5 (orange), R2A (purple), and blb1 (green). Group differences were assessed using two-sided Dunnett-type multiple-comparison tests applied to estimated marginal means (EMMs). Significant differences between transgenic lines and the control are indicated by asterisks above each boxplot: *p < 0.05, **p < 0.01, ***p < 0.001.

**Fig. S8.**
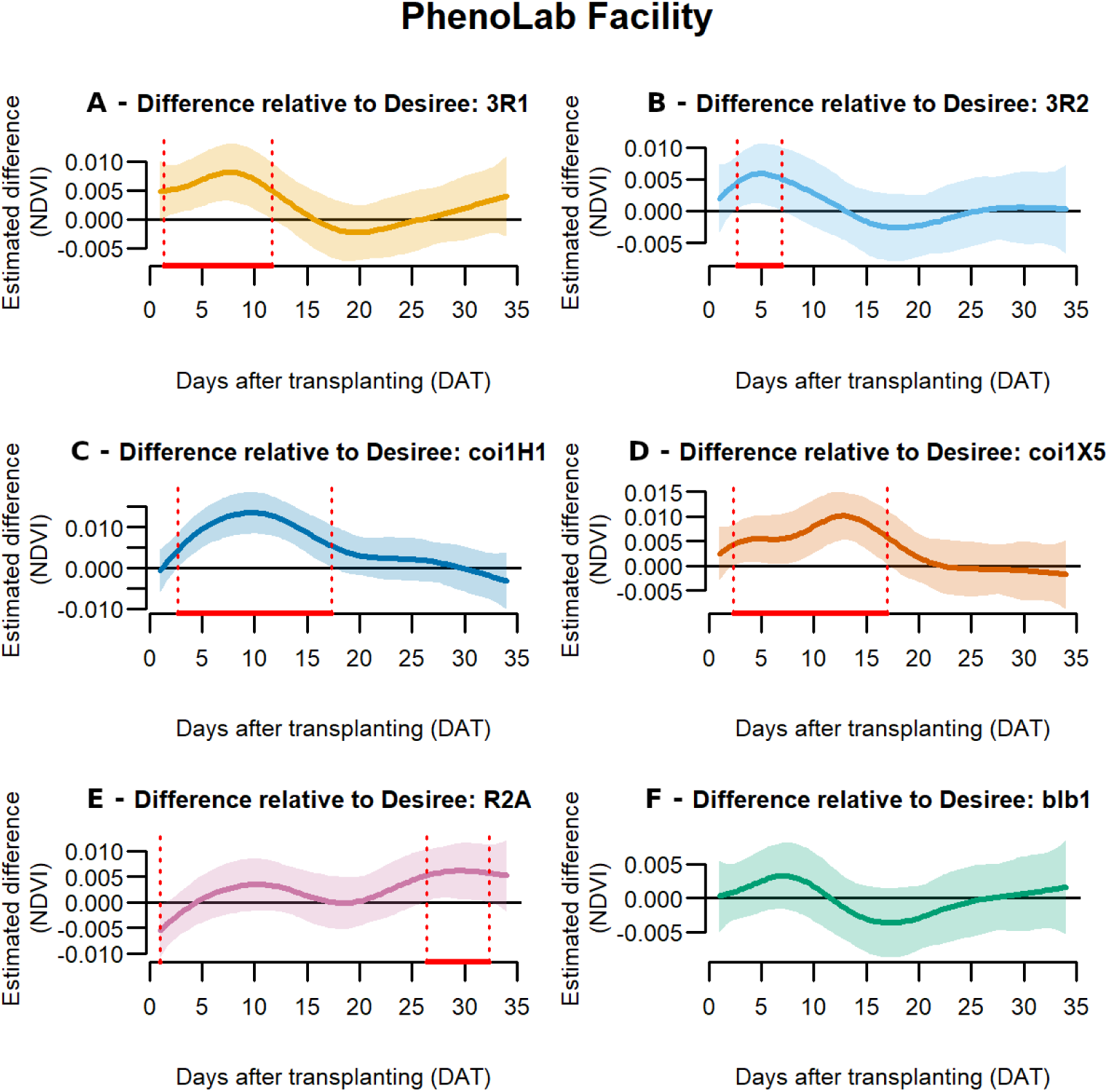
NDVI dynamics in PhenoLab for transgenic lines containing stacked Rpi genes (3R1, 3R2), modified phytohormone pathways (coi1H1, coi1X5), and single Rpi genes (R2A, blb1). Pointwise differences between the NDVI of Desiree (cv.) and the transgenic lines 3R1 (A), 3R2 (B), coi1H1 (C), coi1X5 (D), R2A (E), and blb1 (F) are shown across days after transplanting (DAT). Group differences were evaluated using two-sided pairwise smooth-difference tests derived from the generalized additive mixed models (GAMMs), with significance concluded when the 95% confidence interval of the difference did not include zero. Six biological replicates per genotype were used. Control plants are shown in grey; transgenic lines are color-coded as follows: 3R1 (yellow), 3R2 (pale blue), coi1H1 (dark blue), coi1X5 (orange), R2A (purple), and blb1 (green). Red lines at the bottom of the panels indicate periods during which significant differences at the 95% confidence level were detected.

**Fig. S9.**
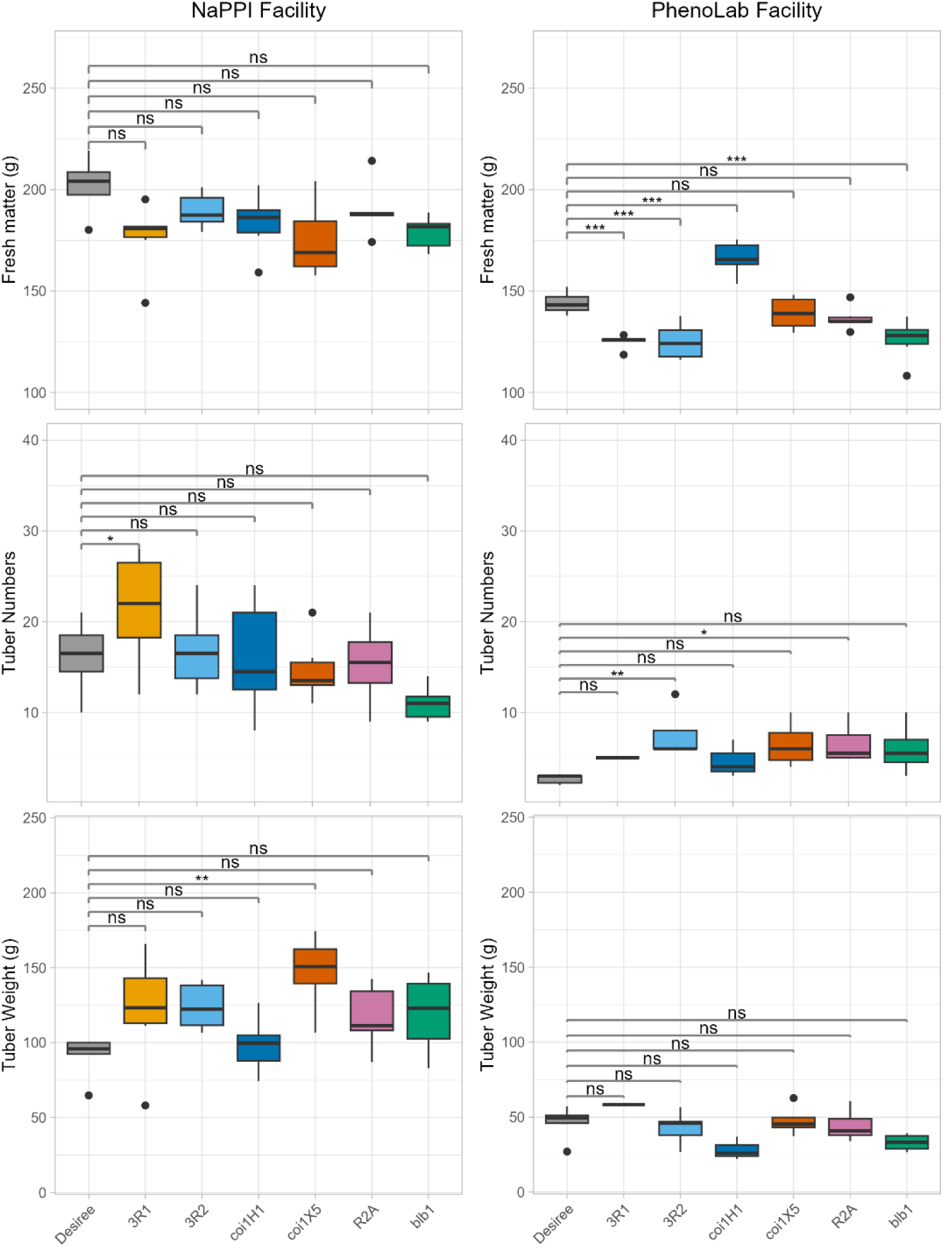
Yield parameters response in transgenic lines containing stacked Rpi genes (3R1, 3R2), modified phytohormone pathways (coi1H1, coi1X5), and single Rpi genes (R2A, blb1). Boxplots showing above-ground fresh weight (top panels), number of tubers (middle panels), and total tuber weight (lower panels) for plants grown in NaPPI (left panels) and PhenoLab (right panels) at the end of the experiment. Each genotype includes six biological replicates. Group differences were assessed using two-sided Dunnett-type multiple-comparison tests applied to estimated marginal means (EMMs). Significant differences between IR treatments and the control are indicated by asterisks: *p < 0.05, **p < 0.01, ***p < 0.001.

## Acknowledgements

We thank Aastha Subedi, Helena Mattisson and Mia Mogren for laboratory assistance at SLU Alnarp; Daniel Richterich and René Hvidberg Petersen for support with plant growth at NaPPI and PhenoLab, respectively; and Valentina Rossi, Svante Resjö, and Junfeng Gao for sampling assistance.

## Author Contributions

**Sylvain Poque:** Investigation, Formal analysis, data curation, image analysis, visualisation & writing. **Murilo Sandroni**: Methodology, Investigation, Formal analysis, Writing & Visualization. **Pedro Caparros:** sample processing and enzyme activity assays. **Jesper Cairo Westergaard:** Data curation, image analysis, Writing - Review & Editing. **Manzur E Mohsina Ferdous:** enzyme activity assays, biochemical data evaluations and quality control. **Katriina Mouhu**: Phenotyping experimentation, Software Analysis, Data Curation. **Erik Andreasson**: Conceptualization, Methodology **Laura Grenville-Briggs Didymus**: Conceptualization, Supervision, Writing - Review & Editing. **Åsa Lankinen**: Conceptualization, Funding acquisition, Supervision, Writing - Review & Editing. **Thomas Georg Roitsch:** Conceptualization, Methodology. **Kristiina Himanen**: Resources, Writing - Review & Editing. **Erik Alexandersson**: Conceptualization, Supervision, Resources, Funding acquisition, Writing, Review & Editing.

## Data availability

Phenomics data will be uploaded on PHIS information system to allow open access. Both raw and processed image data and excel data sheets will be provided.

## References

Abreha KB, Alexandersson E, Vossen JH, Anderson P, Andreasson E. 2015. Inoculation of Transgenic Resistant Potato by Phytophthora infestans Affects Host Plant Choice of a Generalist Moth. PLOS ONE 10, e0129815.

Alexandersson E, Keinänen M, Chawade A, Himanen K. 2018. Nordic research infrastructures for plant phenotyping. Agricultural and Food Science 27, 7–16.

Alexandersson E, Mulugeta T, Lankinen Å, Liljeroth E, Andreasson E. 2016. Plant Resistance Inducers against Pathogens in Solanaceae Species-From Molecular Mechanisms to Field Application. International journal of molecular sciences 17.

Amby DB, Westergaard JC, Großkinsky DK, Jensen SM, Svensgaard J, Liu F, Christensen S, Roitsch T. 2025. The PhenoLab – an automated, high-throughput phenotyping platform for analyzing development, abiotic stress responses and pathogen infection in model and crop plants. Smart Agricultural Technology 11, 100845.

Andrivon D, Pilet F, Montarry J, Hafidi M, Corbière R, Achbani EH, Pellé R, Ellissèche D. 2007. Adaptation of *Phytophthora infestans* to Partial Resistance in Potato: Evidence from French and Moroccan Populations. Phytopathology® 97, 338–343.

Bates D, Mächler M, Bolker BM, Walker SC. 2015. Fitting Linear Mixed-Effects Models Using lme4. Journal of Statistical Software 67, 1–48.

Bengtsson T, Holefors A, Witzell J, Andreasson E, Liljeroth E. 2014a. Activation of defence responses to Phytophthora infestans in potato by BABA. Plant Pathology 63, 193–202.

Bengtsson T, Weighill D, Proux-Wéra E, et al. 2014b. Proteomics and transcriptomics of the BABA-induced resistance response in potato using a novel functional annotation approach. BMC Genomics 15, 315.

Berger S, Sinha AK, Roitsch T. 2007. Plant physiology meets phytopathology: plant primary metabolism and plant-pathogen interactions. Journal of experimental botany 58, 4019–4026.

Bigeard J, Colcombet J, Hirt H. 2015. Signaling mechanisms in pattern-triggered immunity (PTI). Molecular Plant 8, 521–539.

Bradski G. 2000. The OpenCV Library. Dr. Dobb’s Journal of Software Tools in press.

Brown JKM, Rant JC. 2013. Fitness costs and trade-offs of disease resistance and their consequences for breeding arable crops. Plant Pathology 62, 83–95.

Burra DD, Berkowitz O, Hedley PE, Morris J, Resjö S, Levander F, Liljeroth E, Andreasson E, Alexandersson E. 2014. Phosphite-induced changes of the transcriptome and secretome in Solanum tuberosum leading to resistance against Phytophthora infestans. BMC plant biology 14.

Byarugaba AA, Baguma G, Jjemba DM, Faith AK, Wasukira A, Magembe E, Njoroge A, Barekye A, Ghislain M. 2021. Comparative Phenotypic and Agronomic Assessment of Transgenic Potato with 3 R-Gene Stack with Complete Resistance to Late Blight Disease. Biology 10.

Calvo P, Nelson L, Kloepper JW. 2014. Agricultural uses of plant biostimulants. Plant and Soil 383, 3–41.

Cipollini D, Walters D, Voelckel C. 2017. Costs of Resistance in Plants: From Theory to Evidence. Annual Plant Reviews online 47, 263–307.

Cohen Y, Vaknin M, Mauch-Mani B. 2016. BABA-induced resistance: milestones along a 55-year journey. doi: 10.1007/s12600-016-0546-x.

von Dahlen JK, Schulz K, Nicolai J, Rose LE. 2023. Global expression patterns of R-genes in tomato and potato. Frontiers in Plant Science 14.

De Diego N, Spíchal L. 2022. Presence and future of plant phenotyping approaches in biostimulant research and development. Journal of Experimental Botany 73, 5199–5212.

Ehness R, Ecker M, Godt DE, Roitsch T. 1997. Glucose and Stress Independently Regulate Source and Sink Metabolism and Defense Mechanisms via Signal Transduction Pathways Involving Protein Phosphorylation. The Plant cell 9, 1825–1841.

Fimognari L, Dölker R, Kaselyte G, Jensen CNG, Akhtar SS, Großkinsky DK, Roitsch T. 2020. Simple semi-high throughput determination of activity signatures of key antioxidant enzymes for physiological phenotyping. Plant methods 16.

Garcia-Ruiz H, Szurek B, Van den Ackerveken G. 2021. Stop helping pathogens: engineering plant susceptibility genes for durable resistance. Current Opinion in Biotechnology 70, 187–195.

Ghislain M, Byarugaba AA, Magembe E, et al. 2019. Stacking three late blight resistance genes from wild species directly into African highland potato varieties confers complete field resistance to local blight races. Plant Biotechnology Journal 17, 1119–1129.

Großkinsky DK, Svensgaard J, Christensen S, Roitsch T. 2015. Plant phenomics and the need for physiological phenotyping across scales to narrow the genotype-to-phenotype knowledge gap. Journal of experimental botany 66, 5429–5440.

Hadley Wickham. 2016. ggplot2: Elegant Graphics for Data Analysis. Journal of the Royal Statistical Society Series A: Statistics in Society 174, 245–246.

Haitz M, Lichtenthaler HK. 1988. The Measurement of Rfd-Values as Plant Vitality Indices with the Portable Field Chlorophyll Fluorometer and the Pam-Fluorometer. Applications of Chlorophyll Fluorescence in Photosynthesis Research, Stress Physiology, Hydrobiology and Remote Sensing, 249–254.

Halim VA, Altmann S, Ellinger D, Eschen-Lippold L, Miersch O, Scheel D, Rosahl S. 2009. PAMP-induced defense responses in potato require both salicylic acid and jasmonic acid. Plant Journal 57, 230–242.

He Z, Webster S, He SY. 2022. Growth–defense trade-offs in plants. Current Biology 32, R634–R639.

Van Hulten M, Pelser M, Van Loon LC, Pieterse CMJ, Ton J. 2006. Costs and benefits of priming for defense in Arabidopsis. Proceedings of the National Academy of Sciences of the United States of America 103, 5602–5607.

Hunter JD. 2007. Matplotlib: A 2D graphics environment. Computing in Science and Engineering 9, 90–95.

Jammer A, Akhtar SS, Amby DB, Pandey C, Mekureyaw MF, Bak F, Roth PM, Roitsch T. 2022. Enzyme activity profiling for physiological phenotyping within functional phenomics: plant growth and stress responses. Journal of Experimental Botany 73, 5170–5198.

Jammer A, Gasperl A, Luschin-Ebengreuth N, et al. 2015. Simple and robust determination of the activity signature of key carbohydrate metabolism enzymes for physiological phenotyping in model and crop plants. Journal of experimental botany 66, 5531–5542.

Kalaji HM, Jajoo A, Oukarroum A, Brestic M, Zivcak M, Samborska IA, Cetner MD, Łukasik I, Goltsev V, Ladle RJ. 2016. Chlorophyll a fluorescence as a tool to monitor physiological status of plants under abiotic stress conditions. Acta Physiologiae Plantarum 2016 38:4 38, 102-.

Karasov TL, Chae E, Herman JJ, Bergelson J. 2017. Mechanisms to Mitigate the Trade-Off between Growth and Defense. The Plant cell 29, 666–680.

De Kesel J, Conrath U, Flors V, et al. 2021. The Induced Resistance Lexicon: Do’s and Don’ts. Trends in Plant Science 26, 685–691.

Lankinen P, Abreha KB, Alexandersson E, Andersson S, Andreasson E. 2016. Genetics and Resistance Nongenetic Inheritance of Induced Resistance in a Wild Annual PlantÅsa. doi: 10.1094/PHYTO-10-15-0278-R.

Lenman M, Ali A, Mühlenbock P, Carlson-Nilsson U, Liljeroth E, Champouret N, Vleeshouwers VGAA, Andreasson E. 2016. Effector-driven marker development and cloning of resistance genes against Phytophthora infestans in potato breeding clone SW93-1015. TAG. Theoretical and applied genetics. Theoretische und angewandte Genetik 129, 105–115.

Lenth R V., Piaskowski J. 2017. emmeans: Estimated Marginal Means, aka Least-Squares Means. CRAN: Contributed Packages doi: 10.32614/CRAN.package.emmeans.

Lichtenthaler HK, Buschmann C, Knapp M. 2005. How to correctly determine the different chlorophyll fluorescence parameters and the chlorophyll fluorescence decrease ratio RFd of leaves with the PAM fluorometer. Photosynthetica 2005 43:3 43, 379–393.

Liljeroth E, Bengtsson T, Wiik L, Andreasson E. 2010. Induced resistance in potato to Phytphthora infestans – effects of BABA in greenhouse and field tests with different potato varieties. European Journal of Plant Pathology 127, 171–183.

Liljeroth E, Lankinen Å, Wiik L, Burra DD, Alexandersson E, Andreasson E. 2016. Potassium phosphite combined with reduced doses of fungicides provides efficient protection against potato late blight in large-scale field trials. Crop Protection 86, 42–55.

Luna E, Van Hulten M, Zhang Y, et al. 2014. Plant perception of β-aminobutyric acid is mediated by an aspartyl-tRNA synthetase. Nature chemical biology 10, 450–456.

McKinney W. 2010. Data Structures for Statistical Computing in Python. SciPy 2010, 56–61.

Millet EJ, Rodriguez Alvarez MX, Perez Valencia DM, Sanchez I, Hilgert N, van Rossum B-J, van Eeuwijk F, Boer M. 2025. statgenHTP: High Throughput Phenotyping (HTP) Data Analysis. CRAN: Contributed Packages doi: 10.32614/CRAN.package.statgenHTP.

Mohammed U, Davis J, Rossall S, Swarup K, Czyzewicz N, Bhosale R, Foulkes J, Murchie EH, Swarup R. 2022. Phosphite treatment can improve root biomass and nutrition use efficiency in wheat. Frontiers in Plant Science 13, 1017048.

Mulugeta T, Abreha K, Tekie H, Mulatu B, Yesuf M, Andreasson E, Liljeroth E, Alexandersson E. 2019. Phosphite protects against potato and tomato late blight in tropical climates and has varying toxicity depending on the Phytophthora infestans isolate. Crop Protection 121, 139–146.

Ngou BPM, Jones JDG, Ding P. 2022. Plant immune networks. Trends in Plant Science 27, 255–273.

Noir S, Bömer M, Takahashi N, Ishida T, Tsui TL, Balbi V, Shanahan H, Sugimoto K, Devoto A. 2013. Jasmonate Controls Leaf Growth by Repressing Cell Proliferation and the Onset of Endoreduplication while Maintaining a Potential Stand-By Mode. Plant Physiology 161, 1930–1951.

Pandey C, Großkinsky DK, Westergaard JC, Jørgensen HJL, Svensgaard J, Christensen S, Schulz A, Roitsch T. 2021. Identification of a bio-signature for barley resistance against Pyrenophora teres infection based on physiological, molecular and sensor-based phenotyping. Plant Science 313.

Pandey C, Sommer SG, Roitsch T, Schulz A. 2025. Phenotyping-based spectral signatures uncover barley cultivars’ sensitivity to combined mildew and drought treatment. Smart Agricultural Technology 11, 101000.

Pedregosa F, Varoquaux G, Gramfort A, et al. 2012. Scikit-learn: Machine Learning in Python. Journal of Machine Learning Research 12, 2825–2830.

Pérez-Bueno ML, Pineda M, Barón M. 2019. Phenotyping Plant Responses to Biotic Stress by Chlorophyll Fluorescence Imaging. Frontiers in Plant Science 10, 477268.

Pérez-Zavala FG, Ojeda-Rivera JO, Herrera-Estrella L, López-Arredondo D. 2024. Beneficial Effects of Phosphite in Arabidopsis thaliana Mediated by Activation of ABA, SA, and JA Biosynthesis and Signaling Pathways. Plants 13, 1873.

Ravnikar M, Vilhar B, Gogala N. 1992. Stimulatory effects of jasmonic acid on potato stem node and protoplast culture. Journal of Plant Growth Regulation 1992 11:1 11, 29–33.

van Rij J, Wieling M, Harald Baayen R, van Rijn H. 2025. itsadug: Interpreting Time Series and Autocorrelated Data Using GAMMs. in press.

Sandroni M, Liljeroth E, Mulugeta T, Alexandersson E. 2020. Plant resistance inducers (PRIs): Perspectives for future disease management in the field. CAB Reviews: Perspectives in Agriculture, Veterinary Science, Nutrition and Natural Resources 15.

Signorelli S, Casaretto E, Etchemendy-Gamundi M, Bentancor M, Harvey Millar A. 2025. The Green Index: A Widely Accessible Method to Quantify the Degree of Greenness of Photosynthetic Organisms. Plant, Cell and Environment 48, 8027–8043.

Spanic V, Vukovic A, Cseplo M, Vukovic R, Buchvaldt Amby D, Cairo Westergaard J, Puskas K, Roitsch T. 2023. Early leaf responses of cell physiological and sensor-based signatures reflect susceptibility of wheat seedlings to infection by leaf rust. Physiologia plantarum 175.

Trejo-Téllez LI, Gómez-Merino FC. 2018. Potassium phosphite (KPhi): A multifunctional fertilizer–pesticide for sustainable agriculture. Journal of Plant Nutrition 41, 625–642.

Tschiersch H, Junker A, Meyer RC, Altmann T. 2017. Establishment of integrated protocols for automated high throughput kinetic chlorophyll fluorescence analyses. Plant Methods 2017 13:1 13, 54-.

Vlot AC, Sales JH, Lenk M, Bauer K, Brambilla A, Sommer A, Chen Y, Wenig M, Nayem S. 2021. Systemic propagation of immunity in plants. The New phytologist 229, 1234–1250.

Vos IA, Pieterse CMJ, Van Wees SCM. 2013. Costs and benefits of hormone-regulated plant defences. Plant Pathology 62, 43–55.

Van Der Vossen E, Sikkema A, Te Lintel Hekkert B, Gros J, Stevens P, Muskens M, Wouters D, Pereira A, Stiekema W, Allefs S. 2003. An ancient R gene from the wild potato species Solanum bulbocastanum confers broad-spectrum resistance to Phytophthora infestans in cultivated potato and tomato. The Plant journal: for cell and molecular biology 36, 867–882.

Wang ES, Kieu NP, Lenman M, Andreasson E. 2020. Tissue Culture and Refreshment Techniques for Improvement of Transformation in Local Tetraploid and Diploid Potato with Late Blight Resistance as an Example. Plants (Basel, Switzerland) 9.

Wang J, Song W, Chai J. 2023. Structure, biochemical function, and signaling mechanism of plant NLRs. Molecular Plant 16, 75–95.

Waskom M. 2021. seaborn: statistical data visualization. Journal of Open Source Software 6, 3021.

Wood S. 2000. mgcv: Mixed GAM Computation Vehicle with Automatic Smoothness Estimation. CRAN: Contributed Packages doi: 10.32614/CRAN.package.mgcv.

Wu L, Gao X, Xia F, Joshi J, Borza T, Wang-Pruski G. 2019. Biostimulant and fungicidal effects of phosphite assessed by GC-TOF-MS analysis of potato leaf metabolome. Physiological and Molecular Plant Pathology 106, 49–56.

Yuan M, Ngou BPM, Ding P, Xin XF. 2021. PTI-ETI crosstalk: an integrative view of plant immunity. Current Opinion in Plant Biology 62.

Zhang H, Liu Y, Zhang X, Ji W, Kang Z. 2023. A necessary considering factor for breeding: growth-defense tradeoff in plants. Stress biology 3.

